# Generation of cardiomyocytes from human induced pluripotent stem cells resembling atrial cells with ability to respond to adrenoceptor agonists

**DOI:** 10.1101/2022.07.20.499551

**Authors:** Faizzan S. Ahmad, Yongcheng Jin, Alexander Grassam-Rowe, Yafei Zhou, Meng Yuan, Xuehui Fan, Rui Zhou, Razik Mu-u-min, Christopher O’Shea, Ayman M. Ibrahim, Wajiha Hyder, Yasmine Aguib, Magdi Yacoub, Davor Pavlovic, Yanmin Zhang, Xiaoqiu Tan, Derek A. Terrar, Ming Lei

## Abstract

Cardiovascular disease is the leading cause of global mortality and morbidity. Cardiac dysrhythmias contribute significantly to this disease burden. Atrial fibrillation (AF) is the most common chronic dysrhythmia. Human induced pluripotent stem cell-derived atrial cardiomyocytes (hiPSC-AMs) present an exciting new model for AF but currently fail to reach maturity and so are limited in translational potential currently. We report a new approach using a combination of Gremlin 2 and retinoic acid treatment of human iPSCs for generating cardiomyocytes resembling atrial cells. More than 40% of myocytes generated by this approach showed rod-shaped morphology, expression of cardiomyocyte proteins (including RyR2 receptors, a-actinin-2, F-actin) and typically a striated appearance, all of which were broadly similar to the characteristics of adult atrial myocytes. Isolated myocytes were electrically quiescent until stimulated to fire action potentials with an atrial myocyte profile and an amplitude of approximately 100 mV, arising from a resting potential of approximately −70 mV. Single-cell RNA sequence (scRNASeq) analysis showed a high level of expression of several atrial specific transcripts including *NPPA*, *MYL7*, *HOXA3*, *SLN*, *KCNJ4*, *KCNJ5* and *KCNA5*. Amplitudes of calcium transients recorded from spontaneously beating cultures were increased by the stimulation of α-adrenoceptors (activated by phenylephrine and blocked by prazosin) or β-adrenoceptors (activated by isoproterenol and blocked by CGP20712A). Thus, our new method provides an efficient approach for differentiating human atrial myocytes with mature characteristics from hiPSCs. This preparation will be very useful for studying signalling pathways in human atrial myocytes, and provides a valuable model for investigating atrial fibrillation and drug discovery.

## 1. INTRODUCTION

Atrial fibrillation (AF) is the most frequently encountered arrhythmia in clinical practice and represents a significant disease burden worldwide, with a high prevalence and ability to cause morbidity and mortality in the population, particularly in the elderly [1]. Current treatments of AF have major limitations including limited efficacy and significant adverse effect liability. These limitations have inspired substantial efforts concerning mechanistic research and innovative approaches for the development of new therapies, such as tailoring treatment to the underlying pathophysiology of AF [2]. For achieving such a goal, suitable model systems replicating the adult human atrial cardiomyocyte phenotype are required.

In recent years, human induced-pluripotent stem cells (iPSCs) have emerged as an alternative in vitro model system to the use of animals for investigation of human disease mechanisms and development of new medications. Numerous disease-specific iPSC lines have been produced for modelling congenital cardiac arrhythmia syndromes, including heritable AF [3; 4; 5]. Although iPSCs hold great promise for heart disease research and treatment, there are obvious obstacles to be overcome. For instance, cardiogenic differentiation of iPSCs by existing protocols often produces a heterogeneous mixture of cardiomyocytes of different subtypes, primarily ventricular cardiomyocytes. More importantly, a major problem is the maturation of cardiomyocytes from hiPSCs, since the resultant cells frequently display developmentally immature characteristics which are analogous to fetal cardiomyocytes ([6; 7]. The immaturity of cardiomyocytes derived from hiPSCs makes them less suitable models for studying the most common heart diseases, that typically occur in adulthood, and also diminishes the suitability of such models for drug screening. Over the past few years, great efforts have been made by a number of groups in developing new approaches in generation of hESC- or hiPSC-derived atrial cardiomyocytes [8; 9; 10; 11; 12; 13; 14; 15]. A particular molecule that has attracted attention and is being used for generating hESC- or hiPSC-derived atrial cardiomyocytes is retinoic acid (RA), a metabolite of vitamin A1, that mediates the functions of vitamin A1 required for growth and development. An earlier elegant study reported by Keller’s group [16] determined RA and WNT signaling as key regulators of human stem cell development. Their study thus provides molecular insights that aid in developing strategies for the generation of atrial cardiomyocytes from pluripotent stem cells. Additionally, other groups have inspected Gremlin2 as a pro-atrial differentiation factor and reported promising results [17]. Gremlin 2 is a signalling molecule involved in cardiac development and atrial specific differentiation as cardiac progenitor cells migrate [17; 18; 19]. These studies have thus prompted the efforts in developing protocols for generating hESC- or hiPSC-derived atrial cardiomyocytes. Atrial cardiomyocytes derived by these different approaches showed enhanced expression of atrium-specific genes and reduced expression of ventricle-specific genes, but, these cells still show important differences from mature atrial myocytes, particularly in their electrophysiological characteristics: for example, such cells frequently lack a stable resting potential, or when there is a stable resting potential the level is substantially more positive than that observed in the adult phenotype.

Here we report a new approach that is able to generate hiPSC-derived atrial myocytes (hiPSC-AMs) showing mature characteristics, using a newly-developed differentiation protocol involving a treatment regimen combining RA and Gremlin 2. With our RA- and Gremlin 2-based differentiation protocol, we observed a high proportion of elongated cells, some of which showed a remarkable adult atrial myocyte-like morphology. Action potentials with an atrial-type morphology were recorded following electrical stimulation of quiescent cells, and the cells also showed a remarkable response to adrenergic stimulation that has not been shown in previous reports in RA-guided differentiation protocols, to the authors’ knowledge [11; 12; 13]. Cardiomyocytes differentiated from hiPSCs using our new RA/Gremlin 2 protocol also showed an atrial-like transcriptomic profile with high level of expression of several atrial specific transcripts including *NPPA* (natriuretic peptide A), *MYL7* (myosin light chain 2, atrial isoform), *KCNJ2* (Kir 2.1 subunit of the inward rectifier carrying I_K1_), *KCNJ5* (Kir3.4 subunit for conducting I_KACh_) and *KCNA5* (K_V_1.5 for conducting I_Kur_) as compared with a control group treated with standard conventional commercial cardiomyocyte differentiation protocol. Our new method therefore provides an effective approach for differentiating human atrial myocytes with mature characteristics from iPSCs. These cells are likely to be very useful for studying the presence and function of components of cell signaling pathways in human atrial myocytes and will likely provide a valuable model for modeling AF disease and drug discovery.

## Materials and equipment

### Materials

- mTeSR™1 medium (STEMCELL Technologies cat. no. 85850)
- PSC Cardiomyocyte Differentiation Kit (Gibco™ Technologies, cat. no. A2921201) containing:

♦ Cardiac Differentiation Medium A
♦ Cardiac Differentiation Medium B
♦ Cardiac Maintenance Medium
- Corning® Matrigel® hESC-Qualified Matrix (Corning™, cat. no. 354277)
- DPBS, no calcium, no magnesium (500mL) (Gibco™ Technologies, cat. no. 14190250)
- ReLeSR™ (STEMCELL Technologies, cat. no. 5872)
- ACCUTASE™ (STEMCELL Technologies, cat. no. 7920)
- Retinoic acid (500mg) (Sigma-Aldrich, cat. 302-79-4)
- Dimethyl sulfoxide (Sigma-Aldrich, cat. no. 67-68-5)
- Recombinant Human GREM2 (R&D system, cat. no. 8436-PR)
- Y-27632 (STEMCELL Technologies, cat. no.72304)

### Equipment

- Water bath, 37 °C
- Cell culture centrifuge
- CO 2 incubator
- ‘Mr Frosty’ freezing box
- Glass hemocytometer
- Laminar flow hood
- Liquid nitrogen cell storage tank

Cell culture plastics

- Six-well plates (Falcon, cat. no. 353046)
- Four-well Plates (Nunc, cat. no. 179830))
- µ-Slide 8 Well coverslip (ibidi, cat. no.80826)

### Procedure

#### Culture of hiPSCs ● timing ~1 week

1| Thaw a vial of frozen Matrigel in 4°C Fridge over night. Resuspend the Matrigel in cold mTeSR™1 medium with a dilution factor of 60. Pipette up and down with cold pipette tip. Add 2 mL diluted Matrigel on each well of six-well plate to fully cover culture plate. Store the coated plate at 4°C for one week or, before use, incubate at 37 °C in 5% CO2 for at least 1 hour to solidify Matrigel.
2| After solidification, remove the supernatant Matrigel solution and rinse the coated plate with room temperate DPBS.
3| Partially thaw a vial of hiPSCs from −80°C or liquid nitrogen in a 37 °C water bath by consistently rotating the vial until only a small amount of ice is left.
4| Transfer the partially thawed hiPSCs to 10x volume of mTeSR™1 medium at room temperature.
5| Centrifuge the cells at 200 g or 800 rpm for 4 minutes.
6| Gently remove supernatant and resuspend the cells in 2 mL of mTeSR™1 medium supplemented with 10 μM Y-27632, and then plate the cells on the Matrigel-coated well. Culture for 24 hours at 37 °C and 5% CO_2_. Caution: Harmful if Y-27632 is swallowed, inhaled or in contact with skin. Wear personal protective equipment including protective clothing and gloves when handling the material. Avoid breathing dust/fume/gas/mist/vapors/spray. Use only outdoors or in a well-ventilated area. Wash face, hands and any exposed skin thoroughly after handling.
7| On the next day, replace the medium with fresh mTeSR™1 medium without Y-27632. From this point, continue to incubate the cells, changing the medium every day until hiPSC colonies reach 70% confluency. Depending on the cell line, this can take between 3 and 6 days. (Note: Y-27632 can be used up to 3 days if cell survival is poor).
8| Once hiPSCs reach 70% confluency, passage the hiPSC colonies with ReLeSR™, an undifferentiated iPSC-selective detachment agent as per the following steps. Aspirate culture medium and then wash the cells by room temperature DPBS.
9| Then add 1 ml ReLeSR™ per well and wait 40 seconds. Remove ReLeSR™, leaving only a thin film of fluid and incubate cells at 37 °C incubator for 5 mins.
10| Add 6 mL mTeSR™1 medium on each well of the six-well plate of cells. Tap the plate from the side gently with a finger for 1 minute to facilitate cell detachment. Collect floating hiPSCs by 10 mL stripette and transfer into new Matrigel-coated plates. The iPSC colonies were conventionally passaged at a 1:3 ratio. (NOTE: The incubation time with the thin film of ReLeSR™ can be further reduced if a better selectivity is required.)
11| Continue to incubate the cells for 3–5 days and change the medium every day until the hiPSC colonies reach 70% confluency. **Cardiac differentiation ● timing ~3 weeks**
12| Inspect cultures for spontaneously differentiated colonies and remove them using a P200 pipette. Differentiated colonies typically show loss of defined edges.
13| Coat one well of a 4-well or 24-well cell culture plate with 1:30 diluted Matrigel for 1 hour at 37°C.
14| Dissociate the hiPSC colonies into single cells with Accutase™. To do so, first wash hiPSCs with DPBS.
15| Add 1 mL Accutase™ per well of a six-well plate, and incubate at 37 °C for 5 minutes.
16| Then add 3 mL fresh mTeSR™1 medium per well of a six-well plate to terminate the reaction of Accutase™.
17| Plate the cells on Matrigel-coated 4- or 24-well plate with mTeSR™1 medium supplemented with 10 μM Y-27683.
18| Gently tilt the plate following a forward-backward and left-right pattern to evenly distribute iPSCs. Critical Step: The even distribution of fully separated single iPSCs is critical for successful CM differentiation.
19| Continue to incubate the cells with daily mTeSR™1 medium change until they achieve 80% confluency.
20| When hiPSCs colonies reach 80% confluency, aspirate mTeSR™1 medium and wash with DPBS. Critical Step: The best initial confluency for differentiation may differ across hiPCS lines. The initial confluency for best yield can be measured by parallel differentiating hiPSCs with a gradient of varying confluency from 30% to 100%.
21| Add 0.5 mL Cardiac Differentiation Medium A on each well of 4- or 24-well plate (Note: the day is labelled as Day 0).
22| On day 2, aspirate medium and replace with same volume of Cardiac Differentiation Medium B.
23| On day 4, aspirate medium and replace with same volume of Cardiac Maintenance Medium supplemented with 1 μg/ml Gremlin 2.
24| On day 6, aspirate medium and replace with same volume of Cardiac Maintenance Medium supplemented with 1 μM Retinoic acid. Caution: Harmful if Retinoic acid is swallowed, inhaled or in contact with skin, causing skin irritation and may damaging fertility. Wear personal protective equipment including protective clothing, dust mask type N95, eyeshields and gloves when handling the material. Avoid breathing dust/fume/gas/mist/vapors/spray. Use only outdoors or in a well-ventilated area. Wash face, hands and any exposed skin thoroughly after handling.
25| On day 8, aspirate medium and replace with fresh Cardiac Maintenance Medium supplemented with 1 μM Retinoic acid.
26| From day 10 to day 20, incubate the culture by changing to unsupplemented Cardiac Maintenance Medium every two days.
**(Optional) Further analysis preparation ● timing ~3 days and onwards**
27| From day 20, hiPSC-atrial cardiomyocytes are ready for further analysis.
28| If you wish to examine isolated cells then to do so, dissociate cardiomyocytes by Accutase™.
29| Wash the cardiomyocyte culture by DPBS and replace by Accutase™.
30| Incubate at 37 °C for 8 mins.
31| Add 3 times volume of Cardiac Maintenance Medium to terminate reaction.
32| Pipette cells with 1000 μL tip to further detach them.
33| Transfer and seed on 1:30 Matrigel-coated polymer-made chambered coverslip.
34| Culture cells at 37 °C and 5% CO_2_ incubator for at least two days prior to further assessment.

## Results

### 2.1 Morphological and immunocytochemistry characterization reveals a more mature atrial-like structure in Gremlin 2/RA-treated cells

Previous studies showed that the *in vitro* specification of cardiomyocyte subtypes is initiated after the induction of mesoderm and before the terminal differentiation of cardiac progenitor cells (8). Therefore, we speculated that a combination of the Gremlin 2 and RA treatment within this time window might be more effective for directing differentiation of iPSCs towards a cardiac atrial phenotype. Thus, we developed our atrial-specific differentiation protocol as shown by the schematic diagram in Figure 1A, with the detailed protocol described in Appendix 1. Briefly, in this protocol, Gremlin 2 was added prior to RA. We then characterized the functional and structural properties of the resulting derived cells, as described below. The experiments were conducted from two independent iPSC lines and multiple batches of cells.

**Figure 1.**
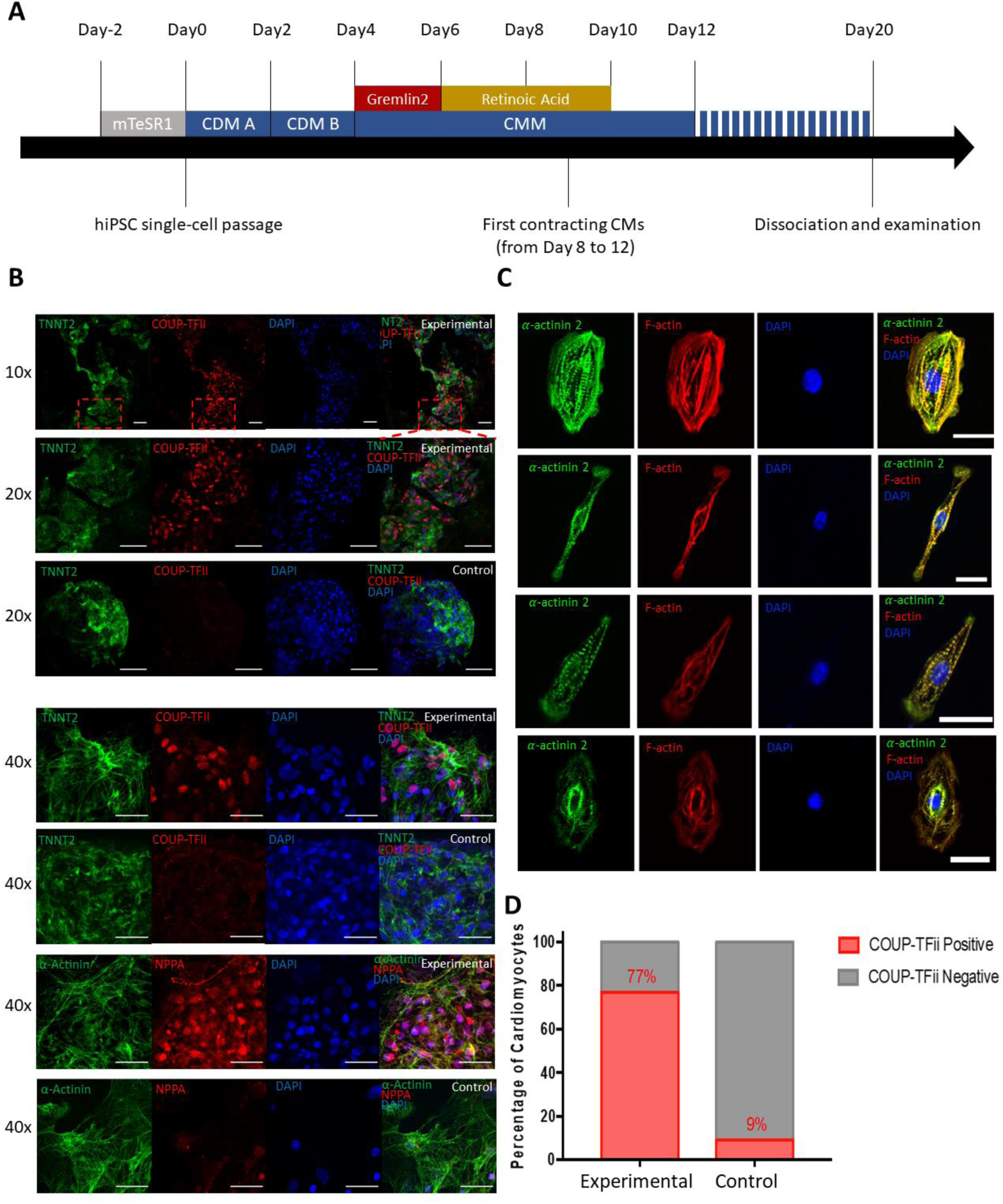
Gremlin2/RA-mediated differentiation of iPSCs into atrial cardiomyocytes. (A) Schematic of the atrial-specific differentiation protocols. Human iPSCs were dissociated into single cardiomyocytes for the differentiation experiments. mTeSR1 is a defined tissue culture medium from Stemcell Technologies. Cardiac Differentiation Medium A (CDM A) including CHIR99021 and Cardiac Differentiation Medium B (CDM B) were applied to both groups. From day 4, all cardiomyocytes were maintained in cardiomyocyte maintenance medium (CMM). In the Experimental group, Gremlin 2 (1 mg/ml) was added at Day 4, and of retinoic acid (1.0 mM) was added with Gremlin 2 still present at Day 6, while the Control group of cells were differentiated without the addition of Gremlin 2 and RA. (B) Functional analysis was performed after day 20 when both cell populations of hiPSC-derived cardiomyocyte showed expression of the cardiomyocytes specific cytoskeleton markers α-actinin 2 (ACTN2) and TNNT2 (cardiac troponin C2), while expression of the atrial cardiomyocytes specific markers COUP-TFII and NPPA, was restricted to the Experimental group. (C) immunofluorescence labelling of the cytoskeleton markers α-actinin 2 and F-actin for cells in the Experimental group demonstrating an elongated morphology with striations. (D) The percentage of COUP-TFII positive cardiomyocytes was 77% in Experimental Gremlin2/RA-treated group. A total of ~1000 DAPI-stained cell nuclear were identified in both treated and control group, and then they were overlapped with COUP-TFII positive nuclear to calculate the percentage of COUP-TFII positive cells. Chi-square test was applied, n=1000, P<0.001.

The cells prepared from the control group (commercial PSC Cardiac Differentiation Kit Gibco™ Technologies, cat. no. A2921201) and cells treated with Gremlin 2/RA (defined as the experimental group) were characterized by immunocytochemistry to determine their morphology and expression of cardiomyocyte-specific proteins, particularly those expressed specifically in atrial myocytes. The cytoskeletal protein ACTN2 (α-actinin 2) and TNNT2 (cardiac troponin C2) in both control and experimental (day 20) cell cultures (Figure 1B). However, labeling for atrial specific NPPA (natriuretic peptide A) and COUP-TFII (also known as nuclear family receptor 2, Group F, member 2) was clearly evident in the experimental group but not control group (Figure 1B). Further immunohistochemistry was carried out in individual cells following single cell isolation from cell cultures (see Materials and Methods). We observed the same general expression of ACTN2 and TNNT2 in both experimental and control isolated cardiomyocyte groups, but only NPPA and COUP-TFII expression in the experimental Gremlin 2/RA-treated group (Figure 1C).

The morphology of some of iPSC-derived atrial myocytes shown in Fig 1C was remarkably similar to that of adult atrial myocytes observed in human and animal hearts [15; 20]. An elongated rod-like morphology resembling the adult phenotype was observed across much of our experimental group (Figure 1C), which was comparable to that reported in human mature atrial myocytes [15; 20]. Furthermore, our Gremlin2/RA experimental group also showed a well-organized striated sarcomeric pattern with sarcomere spacing slightly less than 2 mm (Figure 1C), as observed in adult cardiomyocytes [15; 20]. The percentage of cardiomyocytes in the experimental group that were positive for the atrial-specific COUP-TFII/NPPA was 77% from approximately 1000 counted cells, in contrast to less than 10% in the control group (Figure 1D). Thus, supporting that our hiPSC-AM differentiation protocol established robust protein expression for selected major atrial transcription factors and associated atrial markers.

### 2.2 Transcriptional characterization identifies a more robust atrial transcriptional program in Gremlin 2/RA-treated cells

To gain insights into the molecular signature of the iPSC-derived atrial cardiomyocytes from the experimental group and their differences from the control iPSC-derived cardiomyocytes, we performed single cell RNA sequencing (scRNASeq) on over 300 cells from each control and treatment groups. We first sought to characterize the percentage of atrial cardiomyocytes detected in our sample across an orthogonal gene set to *NR2F2* (COUP-TFII) as in Figure 1. The experimental group showed a high expression of four markers that are characteristic of atrial cardiomyocytes: *HEY1*, *MYL7*, *HOXA3* and *SLN* (Figure 2A). The experimental group was composed 71% of these atrial cardiomyocytes, whereas the control group only 16% (Figure 2A). Thus, our transcriptomic calculation of atrial cardiomyocyte abundance supports our earlier results with proteomic identification and calculation.

**Figure 2.**
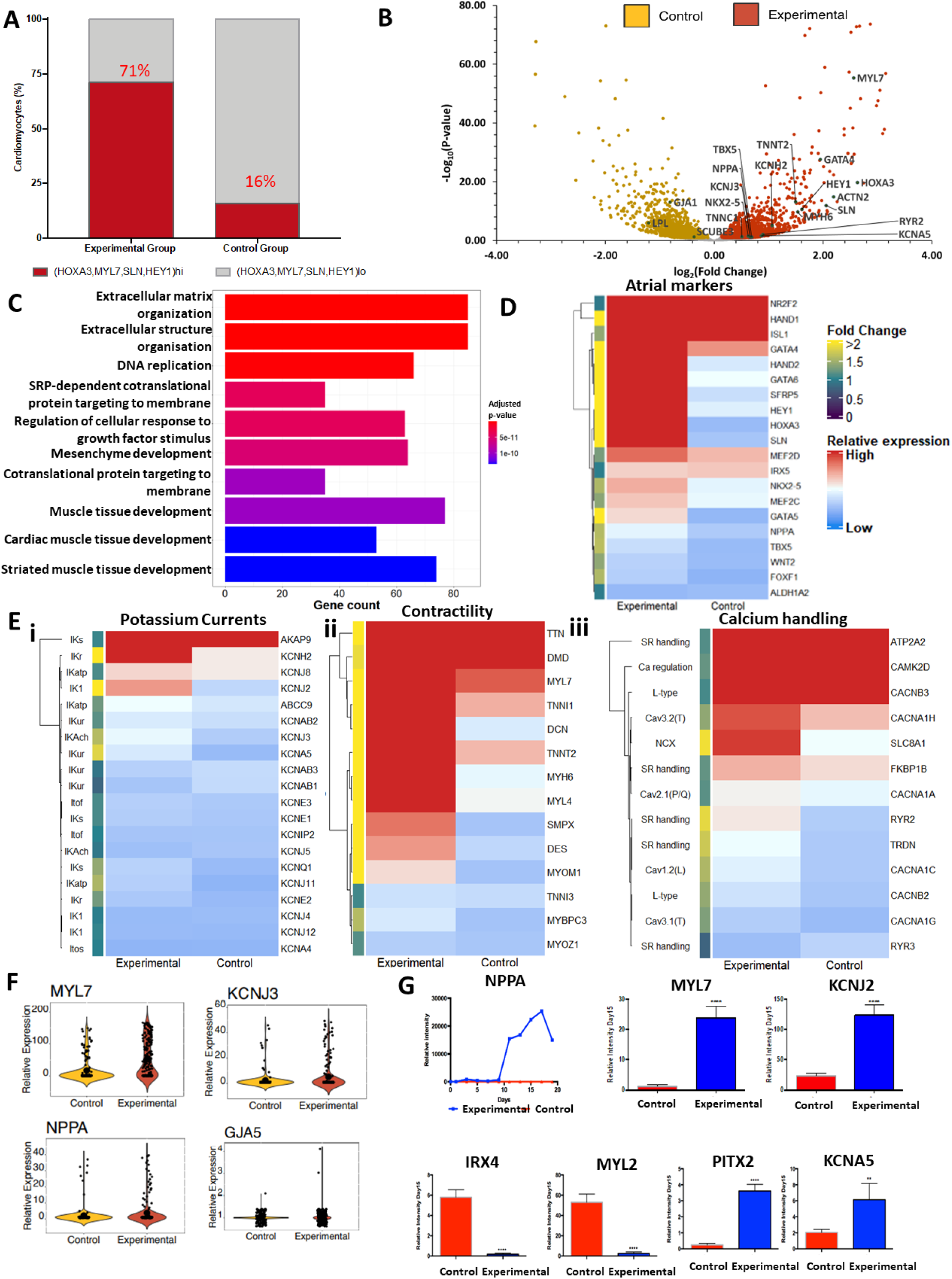
Single cell transcriptomic profile. A) Plot showing the percentage of cardiomyocytes expressing atrial specific markers *HEY1, MYL7, HOXA3 and SLN* in the Experimental and Control groups. B) Differential gene expression analysis showed many up-regulated atrial-specific gene in experimental group with selected markers indicated. C) Gene Ontology (GO) plot showing the top 10 most enriched GO terms in the genes differentially expressed between the control and experimental group. Coloured by Bonferroni-Hochberg-adjusted p-value. Gene count along the x-axis marks the number of genes enriched within each GO term. D) Heatmap plot showing the expression of cardiogenic and chamber-specific markers. Cells are coloured by relative expression. The annotation on the left indicates the specific fold change in expression from the control group to the experimental group. (E) i) Heatmap plot showing the expression of potassium handling genes. Cells are coloured by relative expression. The annotation on the left indicates the specific fold change in expression from the control group to the experimental group. The associated currents for each gene are annotated on the left. ii) Heatmap plot showing the expression of contractility-associated genes. Cells are coloured by relative expression. The annotation on the left indicates the specific fold change in expression from the control group to the experimental group. iii) Heatmap plot showing the expression of calcium handling genes. Cells are coloured by relative expression. The annotation on the left indicates the specific fold change in expression from the control group to the experimental group. The associated currents/roles for each gene are annotated on the left. F) Violin plots of individual expression of atrial genes (*MYL7*, *NPPA*, *KCNJ3, GJA5*) at day 20. G) Time-lapse qRT-PCR gene analysis of *NPPA* over a 20-day time course, and qRT-PCR analysis of expression on day 15 of genes that are specific for either of atrial (*MYL7, KCNJ2, PITX2, KCNA5*) or ventricular (*IRX4, MYL2*) cardiomyocytes. Values shown are relative to housekeeping gene *HPRT*. n=5 batches of cells. Bars show SD of mean. **** indicates P <0.001, and ** indicates P <0.05.

To better understand the differences between our experimental and control groups, we identified the differentially expressed genes between the groups (Figure 2B). Gene expression consistent with atrial cardiomyocytes (including *MLY7, HEY1, HOXA11, SLN, MYH6, KCNH2, GATA4* and *NPPA*) was upregulated in the experimental group, while iPSC-derived cardiomyocytes from the control group showed higher expression of certain ventricular cardiomyocyte-specific genes (including *GJA1* coding for CX43, *LPL* coding for lipoprotein lipase, and *SCUBE3* coding for signal peptide CUB-EG-containing protein 3) (Figure 2B). Notably, we found that genes encoding potassium channels K_V_1.5 (*KCNA5*), and K_V_11.1 (*KCNH2)* displayed significantly higher expression in cardiomyocytes from the experimental group compared with cardiomyocytes from the control group. These potassium channels contribute significantly to the shape and function of APs in human atrial myocytes. *KCNA5* encodes a subunit of a delayed rectifier potassium channel K_V_1.5, underlying the ultra-rapidly activating delayed rectifier K^+^ current I_Kur_ found in human atrial cardiomyocytes [21].

We next sought to further investigate the differentially expressed genes between the two groups beyond the level of isolated genes. We utilized gene ontology (GO) analysis to ascertain if the differentially expressed genes had any enrichment for shared gene function annotations. We identified the top 10 most enriched GO terms, which suggest a difference between groups across a range of significantly enriched functional annotations, relating to organization of the extracellular matrix, replication, and cardiac muscle tissue development (Figure 2C, Supplementary Figures 1–5). Thus, our analysis suggests that our experimental group expresses a diversity of genes of known atrial significance, and that this expression is not just across a few genes but across sets of these genes.

We then sought to identify the expression of specific genes of interest across a range of important modules associated with atrial differentiation and maturation: atrial transcriptional program markers, potassium channels, contractility, and calcium handling. We observe that the experimental group had higher expression of key atrial markers such as *NR2F2, SLN, HEY1, HOXA3,* and *NPPA* (Figure 2D). Whereas, the control group expressed more ventricular markers, such as *IRX5* that is an important driver in development of the ventricular transcriptional program [22] (Figure 2D). However, we note that the experimental group also had increased expression across other genes pertinent for the specification of chamber-specific lineages (Figure 2D).

Potassium currents are particularly important for defining the specific action potential morphology and electrophysiological behavior of atrial cardiomyocytes [23]. We saw an increased expression of several potassium channel-encoding genes (Figure 2Ei). Of particular interest is the higher expression of *KCNJ2*, that encodes K_ir2.1_, which gives rise to the inwardly-rectifying I_K1_ current – a current that demarks more mature atrial cardiomyocytes [24].

Appropriate contractile effort in cardiomyocytes derives from mature molecular organization of actomyosin filaments into sarcomeres connected through to the cell surface and extracellular matrix. Our experimental group has much higher expression of: atrial-specific contractile genes, such as *MYH6, MYL7,* and *MYL4*; more general sarcomeric genes, such as *TTN*, *TNNT2, MYOM1,* and *MYBPC3*; genes that integrate the sarcomeric contraction with the rest of the cytoskeleton and sarcolemma, such as *DMD,* and *DES*; and hypertrophy-associated genes, such as *DCN,* and *SMPX* (Figure 2Eii).

Normal calcium handling is of major importance for normal cardiomyocyte function and the use of hiPSC-derived atrial cardiomyocytes for studying atrial disease. Our data reveals that our experimental group has higher expression of genes associated with both sarcoplasmic reticulum (SR)-associated calcium handling, and membrane calcium handling (Figure 2Eiii). In particular, our experimental group has higher expression of major SR calcium handling genes *RYR2, TRDN, and FKBP1B* alongside its high expression of *ATP2A2* (Figure 2Eiii): whereas the control group had higher expression of the predominantly non-cardiac *RYR3* isoform[25] Additionally, our experimental group has higher expression of *SLC8A1,* which is a key gene for cardiomyocyte ionic homeostasis, and higher expression of T-type calcium channel genes, *CACNA1H* and *CACNA1G* which contribute to atrial-specific electrophysiology, with *CACNA1H* being particularly implicated postnatally[26; 27; 28]

We also demonstrate that our experimental group has higher expression of a range of metabolic genes, including the important transcription factor *PPARGC1A,* which drives mitochondrial biogenesis and oxidative metabolism [29] (Supplementary Figure 6A). Furthermore, we also observe differences in adrenergic signalling-associated genes between our Gremlin 2/RA-treated experimental group and the control, particularly across regulatory cAMP/cGMP phosphodiesterase genes (Supplementary Figure 6B). Finally, we observe robust expression of *GJA1*, and *ATP1A1* across our experimental and control group, and slightly increased *GJA5* expression in our experimental group – suggesting that our experimental group retains expression of expected sodium conductance-related genes, whilst moving towards an atrial phenotype (Supplementary Figure 6C).

We sought to focus on several key genes to further validate these findings. Firstly, we show the expression distribution spread, using our scRNASeq results, showing increased expression across the population of our experimental group for *MYL7, KCNJ3, NPPA,* and *GJA5* (Figure 2F). Then we sought to validate these findings using an orthogonal method: Quantitative PCR (qPCR). *NPPA* expression was examined over the first 20 days, while expression for the other genes of interest were measured on day 15 (Figure 2H). In agreement with our scRNASeq results, qPCR showed significantly higher expression of the atrial-specific genes *NPPA, MLY7,* and preferentially expressed atrial genes such as *KCNJ2, KCNA5,* and *PITX2* in cells from the Gremlin 2/RA treatment group. However, the control group had much higher expression of ventricular-associated genes such as *MYL2* and the transcription factor *IRX4[*[30] Of particular note, the expression of *MYL7*, the gene encoding myosin light chain 2 atrial isoform (*MLC2A*), is 20-fold higher in cells from the experimental Gremlin 2/RA treatment group than in hiPSC-derived cardiomyocytes differentiated by the control commercial protocol, which is consistent with abundance of *MYL7* in adult atrial cardiomyocytes [31]. Therefore, we validated the distinct atrial-like transcriptomic phenotype observed in our Gremlin 2/RA-treated experimental group.

### 2.4. Electrophysiological characterization reveals Gremlin 2/RA treatment produces a more atrial-like electrophysiological behaviour

The electrophysiological properties of atrial cardiomyocytes are distinct from those of ventricular cardiomyocytes. The resting membrane potential for human atrial myocytes has been reported around −74 mV, which is more depolarized than in ventricular cells (c. −81mV) [32]. Additionally, action potentials in atrial cells have a smaller upstroke amplitude with the absence of a prominent plateau phase during the repolarization process [33]. Observations in animal models reproduce these findings that atrial cells have smaller amplitude and shorter duration action potentials [34]. Electrophysiological characteristics were investigated in our Gremlin 2/RA-treated hiPSC-derived atrial cardiomyocytes on day 20-21 of differentiation and compared with the observations in adult human atrial and ventricular myocytes described above. The cultures were dispersed into individual myocytes as described for the immunohistochemistry experiments reported (see Materials and Methods). Action potential recordings of the experimental group cells were obtained by patch clamp under the whole-cell configuration. Patch clamp results revealed an atrial-like action potential in the experimental group cells as shown in Figure 3A. Spontaneous activity was also detected in a small subset of cells (Fig 3A iii and iv) while most others were quiescent until stimulated to fire action potentials by application of current stimuli through the patch electrode (Fig 3A i and ii). Particularly atrial characteristics were the absence of a prolonged plateau and relatively rapid repolarization. The action potential duration at 90 and 50% repolarization (APD_90,_ APD_50_) observed in seven experimental cells were 215 ± 30 ms and 130 ± 30 ms (n=7). The average resting membrane potential of examined hiPSC-derived atrial cardiomyocytes was −67 ± 6 mV (n=7), which is close to that obtained from human atrial cardiomyocytes −74 mV as previously reported [32]. In addition, action potential amplitude (APA) was 99 ± 3 mV recorded in our hiPSC-derived atrial cardiomyocytes. This is comparable to that of human adult atrial cardiomyocytes which is 89 ± 11 mV (n=7). The observations on resting potential, action potential amplitude and duration are shown as bar graphs in Figure 3B.

**Figure 3.**
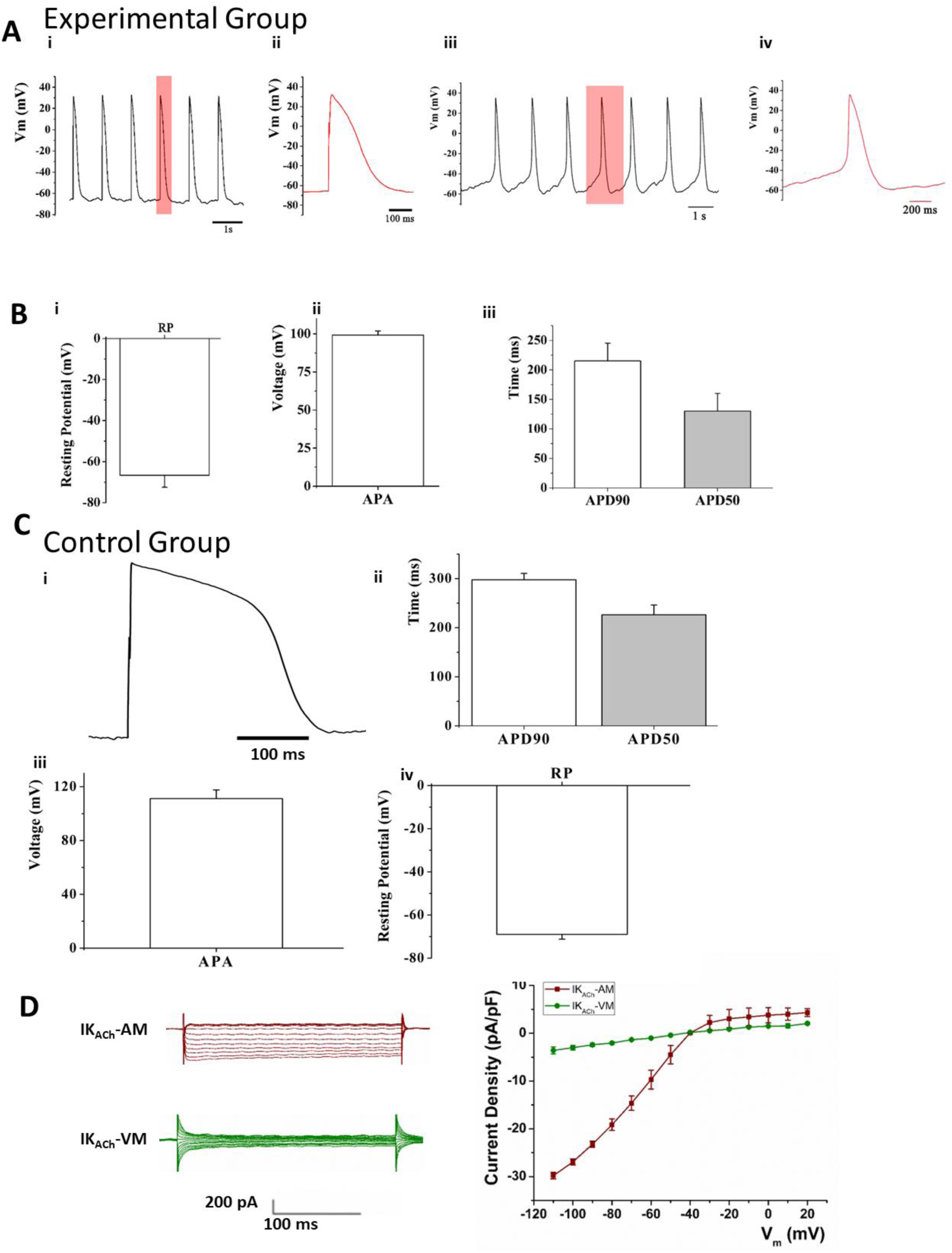
Electrophysiological characterization. A: Action potential traces recorded from quiescent (i, ii) and spontaneous beating (iii, iv) cardiomyocytes in the experimental group with an atrial type morphology. B: Mean values shown as bar graphs for (i) resting membrane potential (RP), (ii) action potential amplitude (APA) and (iii) action potential duration at 90% repolarization (APD90) and 50% repolarization (APD50), measured from seven cells prepared using the experimental protocol. C: Records from cells prepared using the control protocol and having a ventricular phenotype showing (i) a representative action potential trace (ii) Mean APD90 and APD50, (iii) APA and (iv) RP measured from 5 control hiPSC-derived cardiomyocytes. Data are presented as Mean ± SEM. D: Currents induced by exposure to ACh (1 µM) under whole cell voltage clamp conditions for voltage steps in the range −110 to +20 mV recorded from an example cell in the experimental group with an atrial morphology are shown as red traces and labeled as I_KACh_-AM. Corresponding traces from a control cell with a ventricular phenotype showing little or no current induced by ACh are shown in green and labeled as I_KACh_-VM. The IV curve showing a plot of current density against membrane potential (V) is shown in the right panel. The mean currents induced by ACh in cells prepared with the experimental protocol and having an atrial morphology are shown in red. The curve shows the expected inward rectification with a steep increase in current at negative potentials as compared with the flatter slope at depolarized potentials. There was little or no current in cells from the control group with a ventricular phenotype (green)

In contrast, the morphology and characteristics of the action potentials recorded from our control commercially treated cells show ventricular-like features as shown in Figure 3C. For example, the average resting membrane potential of commercially derived hiPSC-cardiomyocytes was −68 ± 2 mV (n=5), The APD_90_ and APD_50_ observed in these same control group cells were 297 ± 13 ms and 226 ± 20 ms (n=5). In addition, APA was 111 ± 6 mV recorded in these control iPSC-CMs (n=5).

To further examine the characteristics of the Gremlin 2/RA-treated hiPSC-derived atrial cardiomyocytes: we assayed their response to acetylcholine (ACh) through the key atrial ACh-activated inwardly rectifying potassium current I_KACh_ ([35] [36]. Figure 3D shows currents induced by exposure to ACh under voltage-clamp conditions. The panel on the right in Figure 3D shows a plot of current against voltage. The ACh-induced current showed the expected inward rectification behavior with a steeper slope of current against voltage for inward currents at more negative potentials as compared to the flatter curve for outward currents at depolarized membrane potentials. In contrast, cells from the control group of hiPSC-CMs, with a more ventricular phenotype, showed little or no current in response to ACh (Figure 3D). The observations showing I_KACh_ in hiPSC-derived atrial cardiomyocytes provide a functional counterpart to the observations in Figure 2 showing robust expression of *KCNJ3* that encodes the protein GIRK1, that in turn comprises the channel which carries I_KACh_ [36].

In summary, the characteristics of action potentials recorded from our Gremlin 2/RA-treated hiPSC-derived atrial cardiomyocytes were broadly similar to those reported for adult human atrial cardiomyocytes and demonstrated differences from the control group consistent with the difference from more ventricular cells.

### 2.5. α- and β-adrenergic stimulation increased the calcium transient amplitude in our Gremlin 2/RA-treated human iPSC-derived atrial cardiomyocytes

We also tested whether our Gremlin 2/RA-treated hiPSC-derived atrial cardiomyocytes responded to adrenoceptor stimulation. The presence of a functional α_1_-adrenoceptor signalling pathway was first tested by exposure of hiPSC-derived atrial cardiomyocytes to phenylephrine (an α-adrenoreceptor agonist), which in adult atrial cardiomyocytes increases Ca^2+^ currents [37]. hiPSC-derived atrial cardiomyocytes from days 25-30 were used to assess the effect of phenylephrine on Ca^2+^ transient amplitude. Figure 4A shows that 10 mM phenylephrine increased Ca^2+^ transient amplitude by 28 ± 10 % as measured from the fluo-4 fluorescence (*P* < 0.05, n = 5 batches of cells) but had no effect on the Ca^2+^ transient rise time or decay time. The involvement of α-adrenoceptors was further evaluated by 1 μM prazosin, a selective a_1_-adrenoceptor antagonist, which by itself had no effect on the amplitude of Ca^2+^ transients (*P* < 0.05, n = 4 batches of cells, data not shown). Figure 4C shows that in the presence of 1 μM prazosin, 10 μM phenylephrine did not cause any significant change in the amplitude of Ca^2+^ transients (*P* > 0.05, n = 4 batches of cells), consistent with blockade of a-adrenoceptors by prazosin and the consequent suppression of the action of the agonist phenylephrine under these conditions (*P* < 0.05, n = 4 batches of cells).

**Figure 4.**
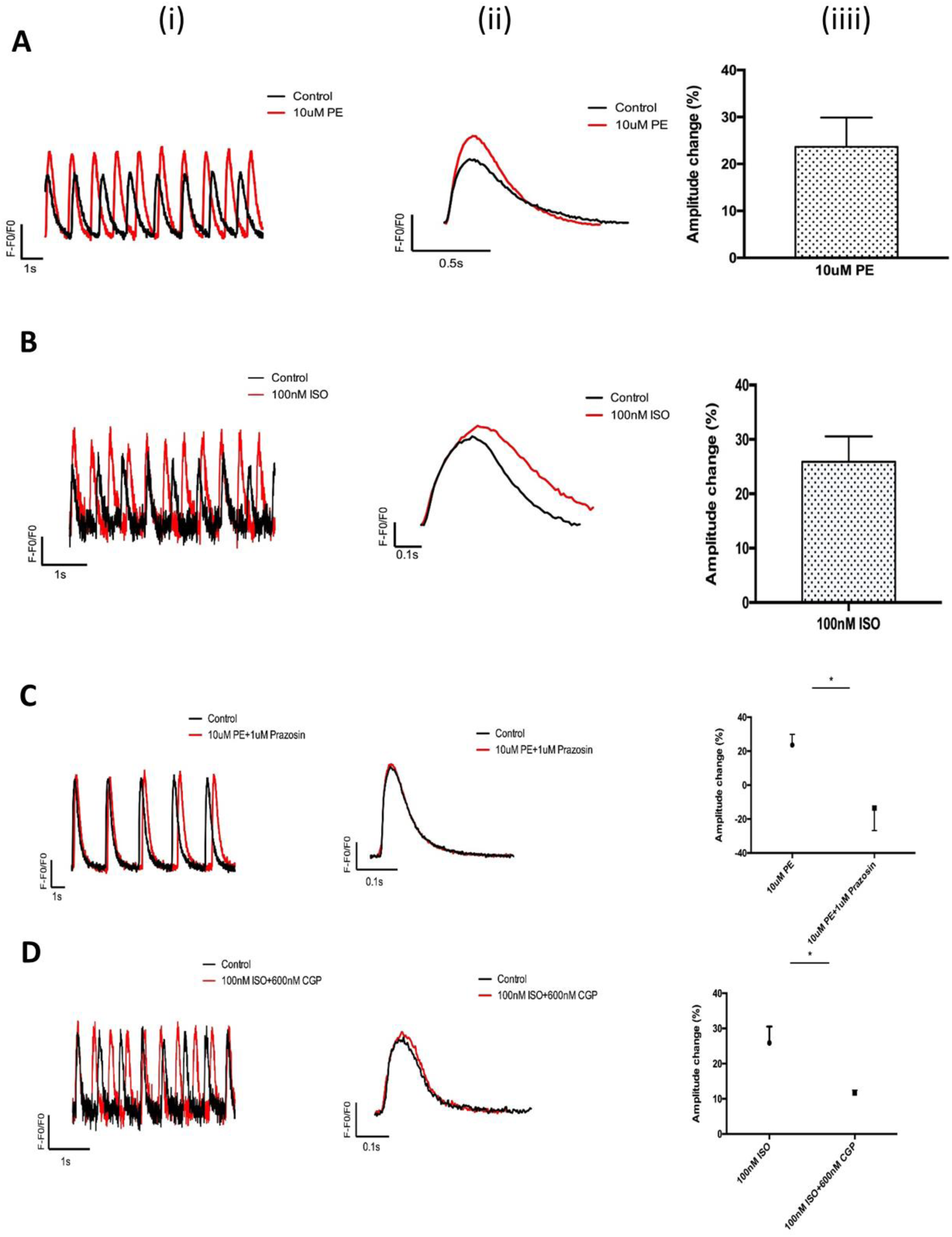
Changes in Ca^2+^ transient (CaT) amplitude in response to α and β adrenoceptor agonists in Experimental group iPSC-CM cultures treated with Gremlin2/RA. (A) CaTs (measured using Fluo-4 as the Ca^2+^ probe) in one representative experiment before and after 10 minutes of exposure to the α-adrenoceptor agonist, phenylephrine (PE, 10 uM). In this and following panels, raw traces are shown in (i), averaged traces are shown in (ii), and graphs representing the mean increase in CaT amplitude are shown in (iii). Red traces in panels (i) and (ii) are in the presence of drugs while black traces are in their absence. A bar graph showing the mean increase in CaT in response to phenylephrine is ahown in (iii). (B) CaTs in a representative experiment before and after 10 minutes of exposure to the β adrenoceptor agonist isoproterenol (ISO, 100 nM). A bar graph showing the mean increase in CaT in response to isoproterenol is ahown in (iii). (C) CaTs in response to PE (10 uM) in the presence of the α-adrenergic antagonist (1 uM prazosin). CaTs after 10 min exposure to PE were abolished by pretreatment with prazosin. Graph showing the mean changes in CaT amplitude in response to PE in the absence and presence of prazosin is shown in (iii). (D) CaTs in response to ISO (100 nM) in the presence of the β1-adrenergic antagonist (300 nM GCP20712A). CaTs after 10 min exposure to ISO were abolished in iPSCs pretreated with GCP20712A. Graph showing the mean changes in CaT amplitude in response to isoproterenol in the absence and presence of GCP20712A is shown in (iii). Four independent experiments were conducted for each drug combination shown in panels A to D (ii) and (iii). Data in A to D (iii) are shown as Mean ± SEM, *P<0.05.

α- and β-adrenergic receptors are highly expressed in human heart, with β-receptors more abundant than α-receptors. β-receptor stimulation leads to increased synthesis of cAMP and activation of PKA which subsequently phosphorylates L-type Ca^2+^ channels and phospholamban, leading to increased Ca^2+^ influx and Ca^2+^ uptake into the SR [Brodde et al., 1999]. As shown in Figure 4B, 10 minutes of 100 nM isoprenaline (a non-selective β agonist) treatment led to an increase of 26 ± 5% in the calcium transient amplitude (*P* < 0.05, n = 4 batches of cells). As shown in Figure 4D, the involvement of β1-adrenoceptors in this response was tested by exposure of the cells to a selective β1 antagonist: CGP20712A (600 nM). When CGP20712A was present, 100 nM isoprenaline had no significant effect on the amplitude of Ca^2+^ transients (*P* > 0.05, n =4 batches of cells). It therefore appears that both α and β adrenoceptors are present and functional in our Gremlin 2/RA-treated hiPSC-derived atrial cardiomyocytes, though we did not detect any changes in calcium removal back in the SR, which were expected.

## DISCUSSION

In the report presented here, we have developed a new protocol to generate functionally responsive atrial cardiomyocytes from human iPSCs. The protocol involves temporal regulation of WNT and BMP signaling and the resultant cells structurally and functionally resemble atrial cardiomyocytes in terms of atrial transcription programs, electrophysiology, and morphology.

The key findings in this report are: i) development and validation of a new protocol consisting of treatment with Gremlin 2 and RA that permits differentiation of cardiomyocytes with an atrial phenotype from human iPSCs; ii) a substantial fraction of cells generated by this protocol showed a rod-shaped morphology with a single central nucleus similar to that observed in cells isolated from human atria; iii) these myocytes demonstrated a distinctly atrial transcriptional program; iv) most isolated rod-shaped cells were electrically quiescent until stimulated to fire typical action potentials with an amplitude of 100 mV arising from a resting potential of approximately −70 mV; v) hiPSC-derived atrial cardiomyocytes from this protocol responded to adrenergic stimuli by both α or β adrenoceptor agonists.

Development of our Gremlin 2 and RA combination treatment protocol for iPSC-atrial myocytes differentiation was based on previous observations showing that both of these agents can separately influence heart development. In zebrafish, Gremlin 2 was essential for the development of the heart [18]. Muller and colleagues also showed that the ventricular and atrial cardiomyocyte specification was affected by the expression of Gremlin 2 because of its role in mediating phosphorylated SMAD1/5/8 and BMP signalling [18]. Additional work [19]showed that Gremlin 2 enhanced cardiac differentiation and increased the atrial cell population in mouse ESC-derived cardiomyocytes, as was evident from electrophysiological studies demonstrating a mixture of different action potential shapes (ventricular, atrial and nodal) in untreated cells, while a dominant atrial-type action potential was recorded from Gremlin 2 treated cells. Gene expression analysis also showed significantly higher expression of atrial-specific genes, *MYL7*, *GJA5*, *KCNJ5*, *CACNA1D1*, *SLN* and *NPPA* in the cells receiving Gremlin 2 treatment [15]. RA has separately been shown to influence heart development, by actions that are thought to involve modification of BMP signalling [38]. Currently, a range of RA-based protocols exist for hiPSC/hESC differentiation into atrial cardiomyocytes and similarly for Gremlin2 [8; 9; 10; 13; 19; 39; 40; 41; 42; 43]. Whilst these differentiation protocols have shown success in establishing an atrial-like phenotype, they mostly produced immature cells, particularly in terms of electrophysiology and pharmacology. Thus, whilst the two agents have been shown separately to reliably induce an atrial-like phenotype, we sought to understand their combinatorial effect. It is interesting to speculate on the precise mechanism through which Gremlin2 and RA synergize, as previously attempts with Noggin and RA have also produced atrial-specific cardiomyocytes, albeit with a less robust atrial phenotype [44]. Both Noggin and Gremlin 2 both possess BMP antagonism[[19]. However, Gremlin2activates *COUP-TFII* and *HEY1* in a JNK-sensitive manner not observed with Noggin signaling [19]. It is beyond the scope of this study, but it will be interesting to further characterize the putative role of JNK signalling in atrial differentiation. The combined use of Gremlin 2 and RA introduced in our experiments described above substantially increased the effectiveness of the new protocol in directing iPSC development towards a more mature atrial phenotype and excitingly across several structural and functional domains of maturity.

An important feature of the cardiac cells produced by the novel methods described here is a morphology that resembles the adult atrial phenotype. The cells had a single nucleus, and approximately 40% showed an elongated ‘rod’ shape that is characteristic of adult myocytes. This was particularly evident in isolated cells that had been dispersed from cell cultures (Figure 1C). Immunohistochemistry demonstrated that expression of atrial-specific proteins, particularly COUP-TFII and NPPA, was greater in cultures from the Gremlin 2/RA-treated experimental group than in cultures from the commercially-derived group (with 78% cardiomyocytes showing COUP-TFII expression in the treated experimental group as compared with 9% in the control group).

A more extensive transcriptome analysis by single-cell RNA sequencing on over 300 cells from each group provided further evidence that Gremlin 2/RA treatment elicited a more atrial transcriptional program than in the control group: atrial genes showing enhanced expression included: *MYL7, HEY1, HOXA3, SLN, MYH6, KCNH2, GATA4 & NPPA*. Whereas, the control group had a predominately more ventricular transcriptomic signature, with greater expression of ventricular transcription factors such as *IRX5* and *IRX4.* We used the unbiased nature of scRNAseq to cross-validate our protein-level expression of atrial markers following Gremlin 2/RA treatment (Figure 1). We observed the Gremlin 2/RA-treatment produced an increased percentage of atrial cardiomyocytes from 16% in the control group to 71% in the experimental group. Particular genes that were upregulated were those that are important for atrial cell function, including ion channels (e.g. *KCNA5, KCNJ2, KCNJ3,* and *KCNH2*) and calcium handling proteins (e.g. *RYR2, CACNA1H, CACNA1G, SLC8A1*). It is interesting to note that our scRNAseq analysis suggests that despite robust expression of *NR2F2* in the control group to a somewhat comparable degree as the experimental group (Figure 2D), this does not appear to be coupled to an increase in the protein levels of its gene product COUP-TFII (Figure 1B,D), nor to an atrial transcriptional program (Figure 2). This highlights the opportunity of in vitro differentiation protocols to study normal developmental signalling and mandates further study to ascertain the epigenetic regulation of *NR2F2*.

In view of the above mRNA expression of genes coding for atrial ion channels, it is perhaps unsurprising that electrophysiological recording provided important additional information supporting the effectiveness of the new Gremlin 2/RA treatment in directing iPSC development towards the adult atrial phenotype. When single cells were isolated from the cultures in which they were grown, they were generally electrically quiescent, and brief current stimuli were needed to initiate action potentials, to the authors’ knowledge this has not been previously reported. The resting membrane potential approached −70 mV, and the amplitude of action potentials was approximately 100 mV consistent with an overshoot of approximately 30 mV. The shape and duration of action potentials was also broadly consistent with the adult atrial phenotype. Similarly, the amplitude of action potentials using the improved protocol with a combination of Gremlin 2 and RA is also larger than in previous reports using either as a differential factor in isolation[9; 42; 44; 45]. In particular, it is important to note the improved activity and larger current we recorded from the preferentially-atrial current I_KACh_ compared to previous RA-only protocols [8; 13], with ours much closer to that seen in adult human atrial cardiomyocytes[46]. Additionally, our robust I_KACh_ signal was concomitant with more clear mature sarcomeric subcellular structure, which was less obvious in other protocols with robust I_KACh_ signal [8]. We highlight I_KAch_ for particular note given the major importance of this current in atrial repolarization and membrane stability and its implication in atrial fibrillation [47; 48] which makes a robust I_KACh_ current likely a necessity for meaningful modeling of atrial fibrillation, drug development, and translational work using hiPSC-derived atrial cardiomyocytes. All these features make cells derived using our new Gremlin 2/RA protocol suitable for the testing of beneficial and harmful effects of drugs on human cardiac atrial muscle.

We also sought to investigate the dominant electrophysiological properties of the two cell culture conditions (experimental against control) beyond individual currents. Whilst it would have been interesting to see the difference between our hiPSC-derived atrial cardiomyocytes, and atrial cardiomyocytes from the control group treated with commercial reagents only, we compared the dominant and modal phenotype in each condition. This ventricular-like and atrial-like behavior from the control and experimental groups, respectively, was representative of all of the cells we assayed in either group via patch-clamp. The modal cell in a given culture will influence the pre-dominant electrophysiological phenotype, and thus our results gives an unbiased consideration of the two groups, without any bias introduced through positive or negative selection of the cells for comparison.

Another important characteristic of the cells that makes them useful experimental models for studying human cardiac muscle function is the ability to respond to the selective agonists: phenylephrine and isoprenaline. The observation that these effects could be blocked by the appropriate antagonists provided further support for the expression and functional intracellular coupling of both α and β adrenoceptors in the cells prepared by the methods described here (Figure 4). The changes in the amplitudes of CaTs that were observed in response to both α and β adrenoceptor stimulation are consistent with the presence and activity of the intracellular cAMP signalling modules downstream of adrenoreceptor stimulation. It was unexpected that we did not detect a clear positive lusitropic effect from β adrenoceptor stimulation, but this point will be the subject of more detailed future studies. From a transcriptomic picture, it is difficult to predict how differences between expression of phosphodiesterases (Supplementary Figure 6B) give rise to functional differences in cAMP signalling but we speculate that there may be further progress to be made in understanding how these cyclic nucleotide domains compare between hiPSC-derived cardiomyocytes and native cardiomyocytes, that might have contributed to our discordancy between what might be accounted for by different cAMP signalling compartmentalization than in native cardiomyocytes. However, to the knowledge of the authors’, ours is the first iPSC-derived atrial cardiomyocyte differentiation protocol to produce a robust response to adrenergic stimuli without further maturation protocols[[8; 9; 10; 11; 12; 13] [14; 15]

### Limitations

Limitations of the study are as follows: It is important to acknowledge that while we believe the morphology of some cells very closely resembles a more mature atrial phenotype it is clear that all current methods based on the use of iPSCs, including use of the protocol described here, still result in significant variability as illustrated in Figure 1C. In the same way, although it is heartening that many isolated atrial cells prepared with our protocol were electrically quiescent, some cells remained spontaneously active. The use of simple whole-cell recording provided useful information on resting potentials and action potential amplitudes, but it is recognised that use of perforated patch avoids problems associated with junction potentials and should be investigated in future experiments. It is again welcome that the cells responded to adrenoceptor agonists (both α and β) and to acetylcholine, but the underlying signalling mechanisms have not been investigated here and will be studied in more detail in future studies, including the use of other agonists such as adenosine. It is also important to acknowledge that there is some batch-to-batch variability in the efficiency of differentiation using protocols for generation of cells from iPSCs, but our use of the protocol described here over a period in excess of two years has consistently provided cells with the more mature atrial phenotype described here. Although we have proposed that the use Gremlin 2 and RA regulates WNT and BNP signalling, this was not investigated in detail and again remains for future study, including the role of other signalling pathways such as JNK. Understanding the underlying differentiation signalling pathways is important for further discovery in this field, but remains beyond the scope of this initial protocol.

In a similar vein as most other monolayer-based hiPSC-derived cardiomyocyte differentiation protocols, we have not examined contractility. This is a common feature not assayed widely across the literature. This probably relates to the difficulty of assaying contraction in monolayers or individual cells without specialized and complex techniques that often have been optimized on isolated native cardiomyocytes [49; 50; 51]outside of the specific hESc/hiPSC-derived cardiomyocyte context. Despite this, novel techniques have arisen to simplify this and apply it to the stem cell-derived cardiomyocyte context [52]. However, we plan in the future to examine these cells in a 3D tissue-like setting to better understand their contractile nature, amongst other assays more convenient in a 3D setting. It will be particularly enlightening to understand the role of our more mature-like atrial cardiomyocyte morphology (Figure 1) on contractile effort and tension. This will be especially interesting given the intriguing signal we see in our scRNASeq results that show upregulation of several markers (*DCN*, *DES*) associated with mature contractile cardiomyocytes, but a less strong signal in others (*TNNI3*) [53; 54; 55]. However, it is also important to note that these markers of maturity have been primarily assays in a mixed, or ventricular-like hiPSC-derived cardiomyocyte context previously, so within the atrial-like context we should be careful about over-reliance on ventricular-like maturation markers.

Our Gremlin 2/RA protocol leaves further ground to push into the adult-like cardiomyocyte level of maturity across all modules of maturity. Techniques for differentiation of hESC/hiPSC-derived cardiomyocytes have been suggested to produce cardiomyocytes resembling the fetal or pre-natal heart [56; 57; 58]. Our hiPSC-derived atrial cardiomyocytes certainly display structure and functional maturity beyond this, as demonstrated by their more mature morphology (Figure 1), atrial-like transcriptional program (Figure 2), stable electrophysiological resting membrane (Figure 3), robust mature atrial-like currents (Figure 3), and a response to adrenergic stimuli (Figure 4). However, it is likely that we are somewhat around the early post-natal stage, rather than a true adult-like maturity. Our hiPSC-AMs display robust expression of a range of maturation-associated markers, such as *DCN*, *DES*, and wider gene sets consistent with more mature functional states (Figure 2). We see a collection of interconnected transcription factors upregulated in our Gremlin 2/RA-treated group (Figure 2). For example, we see *PPARGC1A*, *MEF*, and *GATA* transcription factors upregulated, which in normal development interact in the early postnatal period to drive metabolic maturation and physiological cardiac hypertrophy[59; 60; 61]. However, certain later postnatal transcriptional ‘switches’ do not yet appear to be quite so developed in our scRNAseq data, despite being much more advanced than in our control group (Figure 2). For example, *TNNI3* expression replacing embryonic *TNNI1* has been suggested as a key marker of cardiac maturity[55], and postnatally T-type calcium currents in the atria swap from primarily Ca_V3.2_ (*CACNA1H)*, to Ca_V3.1_ (*CACNA1G)* [27] [62]. Whilst we do see much greater expression of these more mature transcripts than in our control group, the less mature transcripts appear to remain predominant (Figure 2). Thus, whilst we have made progress advancing beyond early embryonic stages to a more peri-/post-natal stage, further maturity will likely require additional maturation techniques.

A further point is that we followed cells for 20 days, and it will be useful in future studies to extend culture periods, for example to 40 days or longer, when we expect cells will show further enhanced maturity. Previous work by others using commercial differentiation kits has suggested that by day 20, although cells display an atrial-like phenotype, that this eventually changes into a more ventricular phenotype[63]. However, we believe that our approach demonstrates a robust cellular commitment and differentiation to an atrial phenotype. Whilst, Burridge et al certainly utilize clever approaches, we believe that our in depth phenotyping of our hiPSC-AMs does not suggest to us that our cells will lose their atrial phenotype. Whilst Burridge et al primarily define atrial phenotype by the relative ratio of MLC2A/MLC2V expression and action potential morphology, we utilize a greater wealth of signals to characterize the atrial phenotype of our cells. Whilst the art of phenotyping has persistent uncertainty across each technique, there has been major speculation about the role of seeding density in action potential morphology from hiPSC-CMs[64]. However, the uncertainty of gene expression markers for phenotyping appears to be primarily focused around difficulty assigning correct spatiotemporal patterning of these [65]. Thus, whilst we also utilize cellular electrophysiology, we sought to reduce our reliance on action potential morphology and utilize alternative electrophysiological and biochemical techniques to characterize our phenotype. We used gene expression markers to phenotype our cell populations (Figures 1–2), showing robust mRNA- and protein-level expression of a range of chamber-specific markers beyond MLC2a/v, including *NR2F2* (COUP-TFII), that acts as a master regulator of atrial identity [31]. However, using single-cell RT-PCR Burridge et al instead show that despite the expression of a number of ‘atrial-like’ genes, their cells at day 20, which appear to be atrial, appear to have a robust expression of *IRX4*, and *MYH7*. *IRX4* is a key transcription factor associated with a ventricular identity [31]. *MYH7* is also associated with a ventricular identity, from mid-to-late development onwards[66]. We also study preferentially atrial currents, such as I_KACh_, to confirm the electrophysiological properties of our cells, beyond action potential morphology. Thus, whilst Burridge et al [63]provide an excellent and intricate warning about the dangers of assumptions about phenotypes, their work focused on non-chamber-specific differentiation, and likely reflects the processes of a heterogenous population. We believe that our protocol shows a robust commitment to an atrial transcriptional program, coupled to an atrial translational program that should maintain an atrial phenotype. However, this remains a clear reminder to all that we should continue to confirm any chamber-specific phenotype across protocols, given the underlying plasticity observed by Burridge et al. The need to confirm the expected phenotype is particularly clear during use of chamber-specific differentiation protocols with non-chamber specific maturation protocols, which we hope to examine in the future.

Our most recent experiments have followed cultures for these longer periods and the cells look broadly similar to those described here, but extensive future work will be necessary to determine whether their morphological, biochemical and electrophysiological characteristics, as well as their ability to respond to different agonists, shows similarities or differences compared with the cells reported in this study. Ultimately, we intend future experiments to ascertain whether our hiPSC-derived atrial cardiomyocytes will be useful for recapitulation of both healthy physiology, and diseased pathophysiology.

In summary, the key features of the cells that mark out advances of the approach presented are: rod shaped morphology in a large fraction of the cells (approximately 40%); expression of atrial-specific transcripts; electrophysiological charactersistics, closely resembling the adult atrial phenoype; and ability to respond to adrenoceptor stimulation. It seems likely that these human cells, with characteristics closely resembling those of the adult atrial phenotype, will provide an important resource for drug testing (particularly drugs to treat atrial fibrillation) and for the investigation of cell signaling mechanisms.

## 1. Experimental procedures

### 1.1. iPSC cell culturing, differentiation and characterizations

The iPSCs lines used for the cardiomyocyte differentiation were purchased from Gibco® Life Technology, Carslbad, USA or developed in the Lab. Cord blood-derived CD34+ progenitor cells were reprogrammed into iPSCs by using a three-plasmid and seven-factor episomal system (OCT4, Sox2, Myc, Klf4, Nanog, SV40LT and Lin28 antigen). iPSC lines were fully characterized by a. immunostaining, b. qPCR, c. tri-lineage differentiation d. teratoma essay e. karyotyping as described in Apendix.

iPSC line was cultured on Matrigel-coated 6-well culture plate with mTeSR™1 medium (STEMCELL™ Technologies), human iPSCs were maintained undifferentiated with daily medium change as the protocol described in details in Apendix.

### 1.2. Cardiomyocyte isolation and immunofluorescent characterization

Cardiomyocyte aggregates were dissociated by using Accutase™ (STEMCELL™ Technologies). Following a DPBS wash, Accutase™ was added to the cardiomyocytes for 8 minutes at 37°C. After the addition of 2-3 volume of fresh CMM to terminate the reaction of Accutase™, the cells were firstly physically detached and collected by 1000uL pipette. Subsequently, dissociated cardiomyocytes were transferred and seeded to 1:30 Matrigel-coated polymer-made coverslip. They were cultured with CMM and 10um Y-27683 for at least two days prior to immunofluorescent examination. The dissociated iPSC-derived cardiomyocytes cultured on the polymer-made coverslips (ibidi Technologies) were washed with PBS and fixed with 4% paraformaldehyde (PFA) at room temperature for 20 minutes and then went through various steps of staining as described in Appendix in details. The immunocytochemistry images were analyzed by ImageJ.

### 5.3 Single cell RNA-sequencing and RT-qPCR

Human iPSC-derived cardiomyocytes were collected as the protocols described in Appendix and were fixed by chilled Methanol and stored on ice for 15 minutes for fixation prior to −80°C storage. The fixed cells were rehydrated using a FACS Aria II or FACSJazz (BD biosciences). Single cells (based on DAPI exclusion and forward/side scatter properties) were sorted into 384-well hard-shell plates (Biorad) with 5 μl of vapor-lock (QIAGEN) containing 100-200 nl of RT primers, dNTPs and synthetic mRNA Spike-Ins and immediately spun down and frozen to −80°C. Sort-sequencing was used for single cell RNAseq. In brief, cardiomyocytes were lysed by 5 min at 65°C, when RT and second strand mixes were dispersed with the Nanodrop II liquid handling platform (GC biotech). Aqueous phase was splited from the oil phase after pooling all cardiomyocytes in one library, followed by IVT transcription. For library preparation, CEL-Seq2 protocol was applied. Primers was composed of a 4bp random molecular barcode, 24 bp polyT stretch, a T7 promoter, a cell-specific 8bp barcode, and the 5′ Illumina TruSeq small RNA kit adaptor. Single-cell mRNA was subsequently reverse transcribed, converted to double-stranded cDNA, assembled and in vitro transcribed for linear as required for the CEL-Seq 2 protocol (Hashimshony et al., 2016). Illumina sequencing libraries were then made with the TruSeq small RNA primers (Illumina) and sequenced paired-end at 75 bp read length the Illumina NextSeq. Read alignment was performed. Subsequently, candidate cells and genes were analyzed in R environment. Cells were filtered using a criteria in which transcript number > 12,000 and gene were filtered > 5 transcript in more than 5 cells. Differential gene expression analysis was performed via RaceID (v0.1.5) [67]]. Violin plots were generated by using R package ggplot2 (v3.2.1) [68] and heatmaps were made by using R package ComplexHeatmap(v2.1.2) [69]]. Gene ontology (GO) enrichment analysis was performed and visualized using clusterProfiler [70].

For RT-qPCR, various gene expressions were analyzed to identify the derived cells as CMs, and to subsequently testify the subtype specificity of the derived CMs. Quantitative PCR analysis was conducted for atrial specific genes (*NPPA*, *MYL7*), inward rectifier potassium channel coding genes (*KCNJ2*), other potassium channel encoding genes (*KCNA5),* and ventricular-specific genes (*MYL2, IRX4).* All the TaqMan® Gene Expression Assays were predesigned by Applied Biosystems by Thermo Fisher Scientific. Quantitative PCR was performed with LightCyclerÒ480 (Roche) using TaqMan® Gene Expression Master Mix (Applied Biosystems by Thermo Fisher Scientific) under the instructions from the manufacturer.

### 5.4 Electrophysiological studies and Ca^2+^ imaging

Action potential recording was recorded by patch clamp under a whole-cell configuration using Axon 700B amplifier system (Molecular Devices, USA). The pipette solution contained (in mmol/L): K-aspartate (K-Asp) 136, KCl 5.4, NaCl 5, MgCl_2_ 1, HEPES 1, EGTA 5, Mg-ATP 5, Phosphocreatine 5, pH 7.2 with KOH. The bath solution consisted of (in mmol/L) NaCl 136, KCl 5.4, MgCl_2_ 1.0, CaCl_2_ 1.8, NaH_2_PO4 0.33, HEPES 5, Glucose 10, pH 7.4 with NaOH. No compensation was applied for changes in junction potential during whole cell recording, but correction with aspartate as the major anion would have resulted in a small shift in recorded potential in the hyperpolarizing direction (30). In experiments where I_KACh_ was to be measured, cells were voltage clamped at a holding potential of −40 mV and step depolarizations were applied in the range −120 mV to +20 mV in increments of 10 mV. Acetylcholine (ACh) was applied at a concentration of 1 µM and currents in the absence of ACh were subtracted from currents in the presence of ACh to give I_KACh_.

Optical mapping experiments were carried out after the cells started to contract. The responsiveness of the derived CMs in the experimental group to β- and α-adrenergic receptor agonists and antagonists was measured. Intracellular calcium transients were analyzed using a 128 x 128 EMCCD camera (Photometrics, Tucson, USA). The derived CMs were pre-loaded with 1 μM Fluo4 (Molecular Probes by Life Technologies) dissolved in DMSO for 15 minutes at 37°C in CMM. Calcium transients of the derived CMs were first recorded without the addition of the drugs as control. Recordings were then taken at 0 minutes, 5 minutes, 10 minutes and 15 minutes after 100 nM isoprenaline (ISO) treatment. Using the same method, the responsiveness of the derived CMs in the experimental group to 10 μM phenylephrine (PE) treatment was tested. In adrenergic receptor antagonist tests, the cells were pre-treated with 600 nM CGP20712A (CGP) (Sigma) dissolved in Fluo4-loading solution prior to 100 nM ISO treatment. Recordings were taken in the same way as described above. Similarly, the cells were pre-treated with 1 μM Prazosin dissolved in Fluo4-loading solution before a treatment of 10 μM PE. Metamorph was used to take the recordings, and 4000 or 8000 frames were taken for each recording, with a frame rate of 100Hz (for 4000 frames) or 333.33Hz (for 8000 frames). Regions of interest were selected using ImageJ. Raw traces were analyzed and baseline corrections were conducted using Clampfit. Calcium transient amplitude change was calculated based on averaged trace. Electrophysiological analysis was performed, in part, using ElectroMap [71] on MATLAB. To calculate beat-to-beat CaT50, the mean signal was segmented into individual beats. Representative regions were defined as two 16×16 pixel regions.

## Data Availability Statement

all data are contained within the manuscript. Reasonable requests for raw data and analysis scripts should be made to the corresponding author.

## Acknowledgements

Our work is supported by the Magdi Yacoub Foundation (FSA, DAT) and British Heart Foundation (BHF) (BHF Centre for Research Excellence (CRE) at Oxford, PG/14/80/31106, PG/16/67/32340, PG/12/21/29473, FS/PhD/20/29053)

## Appendix: expanded information on Experimental Procedures

### 1. Characterization of hiPSC cell lines

The iPSCs lines used for the cardiomyocyte differentiation were purchased from Gibco® Life Technology, Carslbad, USA or developed in the Lab. Cord blood-derived CD34+ progenitor cells were reprogrammed into iPSCs by using a three-plasmid and seven-factor episomal system (OCT4, Sox2, Myc, Klf4, Nanog, SV40LT and Lin28 antigen).

iPSC lines were fully characterized by a. immunostaining, b. qPCR, c. tri-lineage differentiation d. teratoma essay e. karyotyping as follows:

1.1. Immunostaining was used to determine if iPSC express pluripotency markers. iPSC were grown onto irradiated mouse embryonic feeder cells and maintained undifferentiated in serum-free media supplemented with 20% serum replacement (Invitrogen) and 4ng/ml human FGF2 (R&D). Cells were fixed in 4% PFA, cell membranes permeabilized and non-specific staining inhibited by treatment with a blocking solution (Dako). Cells were incubated with primary antibodies against Klf4, Nanog, Oct4, Sox2, SSEA-4 and c-myc, then washed and incubated with the appropriate secondary antibody and DAPI to counterstain the nuclei. For confocal microscopy, slides were analyzed with a Zeiss confocal microscope Laser Scanning Microscope 710 (Carl Zeiss). Pictures were analyzed with Zen 2008 V5,0,0228 software (Carl Zeiss).

1.2. qPCR analysis was used to confirm expression of the endogenous genes Oct4, Sox2, Klf4 and c-myc and simultaneous silencing of the 4 exogenous (retroviral or sendai virus) factors in each of the iPSC clones using published primers (Takahashi, Tanabe et al. 2007).

Cells were collected and total RNA was isolated (Qiagen). cDNA was generated using reverse transcription and qPCR was performed using cybergreen mix (Applied Biosystem), with results normalized to the level of a housekeeping gene, HPRT. Expression of the endogenous genes were compared to gene expression in hESCs, the gold standard of pluripotency. Moreover, we validated that the retrovirus expression for the 4 factors were largely silenced, demonstrating successful reprogramming. Initial analysis confirmed endogenous expression of the pluripotency genes and silencing of the transgenes in two iPSC clones.

1.3. Tri-lineage differentiation in vitro was used to demonstrate functionality. Bona fide pluripotent stem cells are able to give rise to all three germ layers: endoderm, ectoderm and mesoderm. Prior to differentiation, iPSCs were feeder-depleted by passaging onto matrigel.

We used established protocols for tri-lineage differentiation based on two commonly used methods: the formation of three-dimensional aggregates called embryoid bodies (EBs) or culture of cell monolayers on extracellular matrix. To generate EBs, iPSC were dissociated into small clusters of 10-20 cells, and then plated in low attachment plates with the appropriate differentiation media: Endoderm - serum-free media supplemented with 100ng/ml activin A for 5 days. Around D5, we saw that greater than 90% of the cells were c-kit+ CXCR4+ (detected by FACS analysis), demonstrating efficient induction of definitive endoderm. Mesoderm - StemPro-34 media (Invitrogen) supplemented with activin A, BMP4, bFGF, VEGF and the Wnt inhibitor DKK1. Around D6, we visualized detection by FACS analysis the emergence of 3 distinct cell populations, with the c-kitneg KDRlow population signaling the beginning of cardiovascular differentiation. Ectoderm - induction will be done in monolayers in serum-free medium supplemented with Noggin and SB431542 for 11 days. We saw that at D11 greater than 80% of the cells will express Pax6, a marker of neural differentiation.

The teratoma assay is considered the definitive proof of bona fide iPSCs. Tri-lineage differentiation in vivo was obtained by injecting around 1 million iPSC into an immunocompromised mouse, typically sub-cutaneous. Around 6-8 weeks later, teratomas are removed and histological analysis is performed to show the presence of tissues from the 3 germ layers. A human nuclear marker was used to confirm the human nature of the tumor. Karyotype analysis was performed for bona fide iPSC clones. We sent cells to Cell Line Genetics to perform karyotyping using standard protocols.

### 2. iPSC culturing and differentiation

iPSC line was cultured on Matrigel-coated 6-well culture plate with mTeSR™1 medium (STEMCELL™ Technologies), human iPSCs were maintained undifferentiated with daily medium change. mTeSR™1 medium was pre-warmed under room temperature every time before use. The iPSCs were passage every 4 days (mTeSR™1) when cells achieve an approximately 70% confluency. On differentiation day 0, human iPSCs were sufficiently dissociated into single cells for cardiomyocyte differentiation. iPSCs were washed by DPBS and treated by ReLeSR™ for 40 seconds. When ReLeSR™ was aspirated, 1 ml Accutase™ (STEMCELL™ Technologies) was added on the cells and incubated at 37°C for 5 min. After the addition of 2-3 volume of fresh mTeSR™1 medium to terminate the reaction of Accutase™, the cells were plated on 1:30 diluted Matrigel-coated 4- or 24-well plate with mTeSR™1 medium and 10uM Y-27683. The plate was then moved following a forward-backward and left-right pattern to evenly distribute iPSCs. These cells were daily fed with mTeSR™1 medium and 10uM Y-27683 to support single cell proliferation until they achieve 80% confluency. (Note: The even distribution of fully separated single iPSCs is critical for successful CM differentiation)

#### Cardiomyocyte isolation

Cardiomyocyte aggregates were dissociated by using Accutase™ (STEMCELL™ Technologies). Following a DPBS wash, Accutase™ was added to the cardiomyocytes for 8 minutes at 37°C. After the addition of 2-3 volume of fresh CMM to terminate the reaction of Accutase™, the cells were firstly physically detached and collected by 1000uL pipette. Subsequently, dissociated cardiomyocytes were transferred and seeded to 1:30 Matrigel-coated polymer-made coverslip. They were cultured with CMM and 10um Y-27683 for at least two days prior to immunofluorescent examination.

### 3. Immunofluorescence analysis

The dissociated iPSC-derived cardiomyocytes cultured on the conventional glass coverslip or polymer-made coverslips (ibidi Technologies) were washed with PBS and fixed with 4% paraformaldehyde (PFA) at room temperature for 20 minutes. After fixation, the cells were washed 3 times with ice-cold PBS, and permeabilized by 0.1% Triton X-100 in PBS for 20 minutes and wash by PBS three times. Then, they were blocked from non-specific binding by 10% goat serum diluted in 0.1% Tween-20 in PBS (PBST) for 60 min at room temperature. Subsequently, the cardiomyocytes were incubated with primary antibodies diluted in 1% goat serum in PBST in a humidified chamber overnight at 4°C. The dilution for each antibody was listed in the Table 2. After the primary antibody incubation, cardiomyocyte was washed with PBS three times and incubated with secondary antibody diluted in 1% goat serum in PBST for 1.5 hour at room temperature. After the secondary antibody incubation, the cells were washed for three times with PBS at room temperature. All operations post to the involvement of secondary antibody were performed in dark room. Then another glass-made coverslip was transferred onto the ibidi coverslip and mounted with Prolong™ Gold Antifade Reagent Mounting medium with DAPI (Molecular Probes by Life Technologies). Followed by snail polish sealing, the slides can be examined directly or stored in 4°C for approximate one month. The immunocytochemistry images were analyzed by ImageJ.

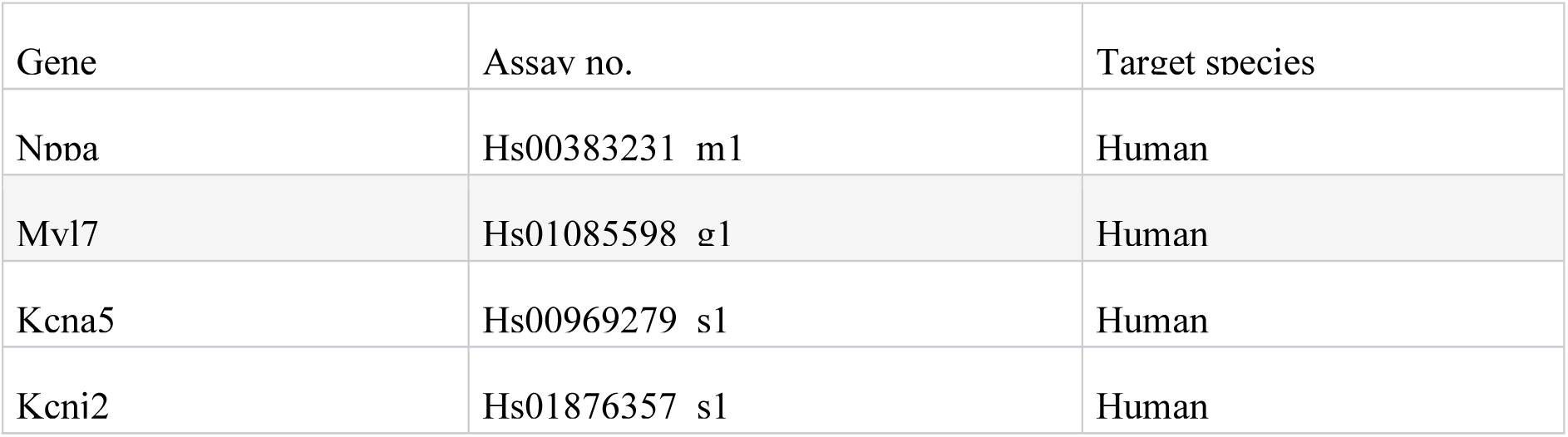
Table of TaqMan® Gene Expression Assays Supplied by Applied Biosystems by Thermo Fisher Scientific

### 5. Single cell RNA-sequencing

#### 5.1. Single-cell dissociation and methanol fixation

Human iPSC-derived cardiomyocytes were briefly washed by DPBS and treated by 300uL pre-warmed 100U/mL Collagenase I (Gibco® Life Technology) dissolved in RPMI 1640 medium (Gibco® Life Technology) for 40 minutes at 37 °C. Then the cells were detached by using 1000uL pipette and return to 37°C for another 10 minuets. Digested cardiomyocytes were then transferred into 1.5mL centrifuge tube containing 1 mL CMM to dampen reaction. After the centrifugation at 300rcf for 3 min using Centrifuge 5424R (Eppendorf), supernatant was discarded when the cardiomyocyte pellet was resuspended in 0.3 mL Accutase (STEM CELL Technologies) and incubated at 37°C for 10 min. Afterward, 1 mL chilled DPBS was applied to the cells to stop the reaction and was subsequently filtered through a 70 mm cell strainer to acquire single cells. The filtered cells were then centrifuged at 300 rcf for 3 mins at 4 °C to reduce cell debris. When supernatant was discarded, the pellet was resuspended by 100uL chilled DPBS, followed by 900uL chilled Methanol and stored on ice for 15 minutes for fixation prior to −80°C storage.

#### 5.2. Single-cell sorting into 384-well hard-shell plate

The fixed cells were rehydrated by applying centrifugation at 3000 rcf for 10min at 4°C, followed by supernatant removal and chilled Rehydration Buffer resuspension (1mL). After using a wide-bored pipette tip to gently pipette mix the cell solution 10 times, the cells were centrifuged, and resuspended following identical process described above to achieve a target cell concentration of 7 x10^5^-1.2 x10^6^/ml). The rehydrated cells were immediately sorted using a FACS Aria II or FACSJazz (BD biosciences). Single cells (based on DAPI exclusion and forward/side scatter properties) were sorted into 384-well hard-shell plates (Biorad) with 5 μl of vapor-lock (QIAGEN) containing 100-200 nl of RT primers, dNTPs and synthetic mRNA Spike-Ins and immediately spun down and frozen to −80°C.

#### 5.3. Sort-sequencing

For SORT-seq, cardiomyocytes were lysed by 5 min at 65°C, when RT and second strand mixes were dispersed with the Nanodrop II liquid handling platform (GC biotech). Aqueous phase was splited from the oil phase after pooling all cardiomyocytes in one library, followed by IVT transcription. For library preparation, CEL-Seq2 protocol was applied. Primers was composed of a 4bp random molecular barcode, 24 bp polyT stretch, a T7 promoter, a cell-specific 8bp barcode, and the 5′ Illumina TruSeq small RNA kit adaptor. Single-cell mRNA was subsequently reverse transcribed, converted to double-stranded cDNA, assembled and in vitro transcribed for linear as required for the CEL-Seq 2 protocol (Hashimshony et al., 2016). Illumina sequencing libraries were then made with the TruSeq small RNA primers (Illumina) and sequenced paired-end at 75 bp read length the Illumina NextSeq.

### 1. RT-qPCR

Various gene expressions were analyzed to identify the derived cells as CMs, and to subsequently testify the subtype specificity of the derived CMs. Quantitative PCR analysis was conducted for early cardiac marker (*NKX2.5*), cardiac troponin coding gene (*TNNT2*), atrial specific genes (*NPPA, PITX2,* and *MYL7*), inward rectifier potassium channel coding genes (*KCNJ2, KCNA5*), and ventricular genes (*IRX4, MYL2*) in experimental and control groups. All the TaqMan® Gene Expression Assays were predesigned by Applied Biosystems by Thermo Fisher Scientific. Quantitative PCR was performed with LightCyclerÒ480 (Roche) using TaqMan® Gene Expression Master Mix (Applied Biosystems by Thermo Fisher Scientific) under the instructions from the manufacturer. In brief, for a total reaction volume of 20 µl, 10 µl of Gene Expression Master Mix, 1 µl of Gene Expression Assays, 1µl of cDNA sample and 8 µl of RNase-free water were used. Cycling parameters were 2 min at 50°C, followed by 10 minutes at 95°C, then followed by 40 cycles of 15 seconds at 95°C and 60 seconds at 60°C. A control for cDNA input was generated by amplifying HPRT1 gene (Applied Biosystems by Life Technologies). The relative gene expression was determined by averaging the results of two or three technical replicates, and comparing the Ct values of genes of interest with those of the control gene using ΔΔCt method.

#### 5.5. Optical Mapping

A custom-designed optical mapping system equipped with an EMCCD camera (Evolve 128, Photometrics, Tucson, AZ, USA) was used. Two green (to excite Ca^2+^ sensitive dye Fluo 4 AM; a peak wavelength of 530 nm) were placed around the imaging chamber so as to uniformly illuminate the sample. CaT measurements were taken at high resolution (512 x 512 pixels; pixel size 16 x 16 μm) at a rate of 1000 Hz. The derived CMs were pre-loaded with 1μM Fluo4 (Molecular Probes by Life Technologies) dissolved in DMSO for 15min at 37°C. Optical mapping experiments were carried out after the cells started to contract. The responsiveness of the derived CMs in the experimental group to β and α adrenergic receptor agonists and antagonists was measured. Calcium transients of the derived CMs were first recorded without the addition of the drugs as control. Recordings were then taken at 0min, 5min, 10min and 15min after 100nM Isoprenaline (ISO) treatment. Using the same method, the responsiveness of the derived CMs in the experimental group to 10μM Phenylephrine (PE) treatment was tested. In adrenergic receptor antagonist tests, the cells were pre-treated with 600nM CGP20712A (CGP) (Sigma) dissolved in Fluo4 loading solution prior to 100nM ISO treatment. Recordings were taken in the same way as described above. Similarly, the cells were pre-treated with 1μM Prazosin dissolved in Fluo4 loading solution before a treatment of 10μM PE. Metamorph was used to take the recordings, and 4000 or 8000 frames were taken for each recording, with a frame rate of 100Hz (for 4000 frames) or 333.33Hz (for 8000 frames). Regions of interest were selected using ImageJ. Raw traces were analyzed and baseline corrections were conducted using Clampfit. Calcium transient amplitude change was calculated based on averaged trace. Electrophysiological analysis was performed, in part, using ElectroMap [71] on MATLAB. To calculate beat-to-beat CaT50, the mean signal was segmented into individual beats. Representative regions were defined as two 16 by 16 pixel regions.

## Supplementary Figure Legends

**Supplementary Figure 1).**
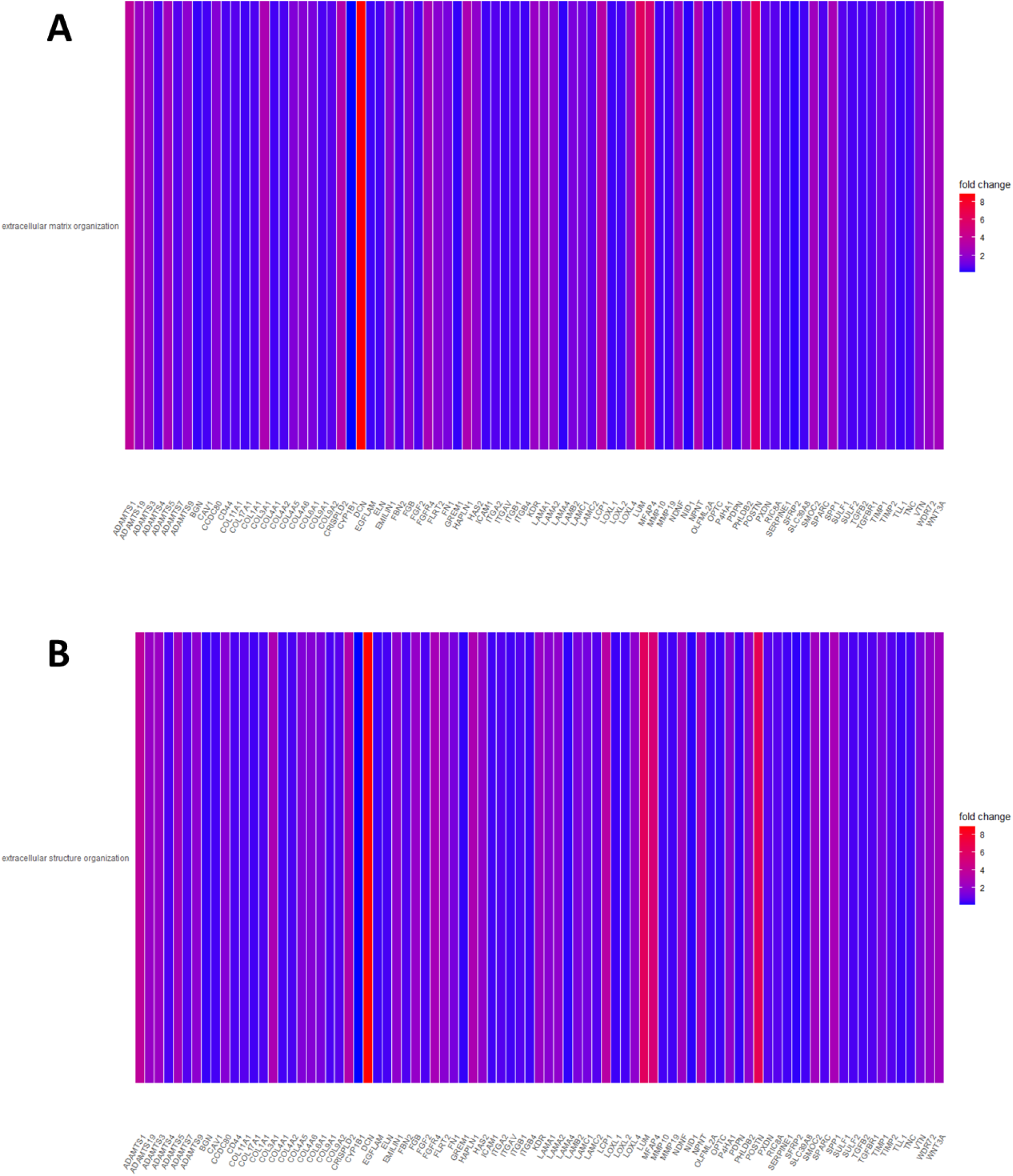
Heatmaps of genes annotated by specific GO terms. A) Heatmap plot showing genes annotated by the GO term: ‘extracellular matrix organisation’. Cells are coloured by fold change of expression from control group to the experimental group. B) Heatmap plot showing genes annotated by the GO term: ‘extracellular structure organisation’. Cells are coloured by fold change of expression from control group to the experimental group.

**Supplementary Figure 2).**
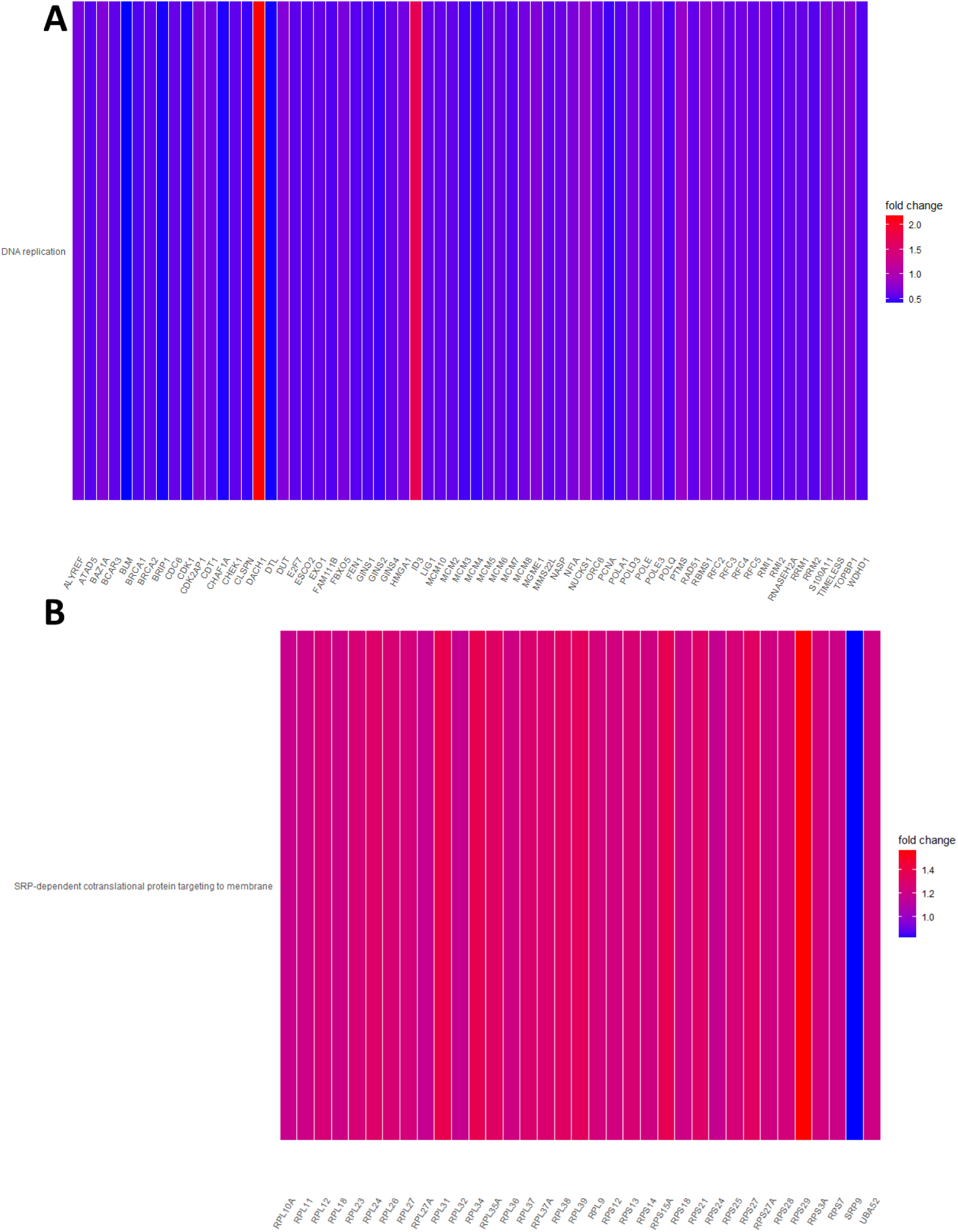
Heatmaps of genes annotated by specific GO terms. A) Heatmap plot showing genes annotated by the GO term: ‘DNA replication’. Cells are coloured by fold change of expression from control group to the experimental group. B) Heatmap plot showing genes annotated by the GO term: ‘SRP-dependent cotranslational protein targeting to membrane’. Cells are coloured by fold change of expression from control group to the experimental group.

**Supplementary Figure 3).**
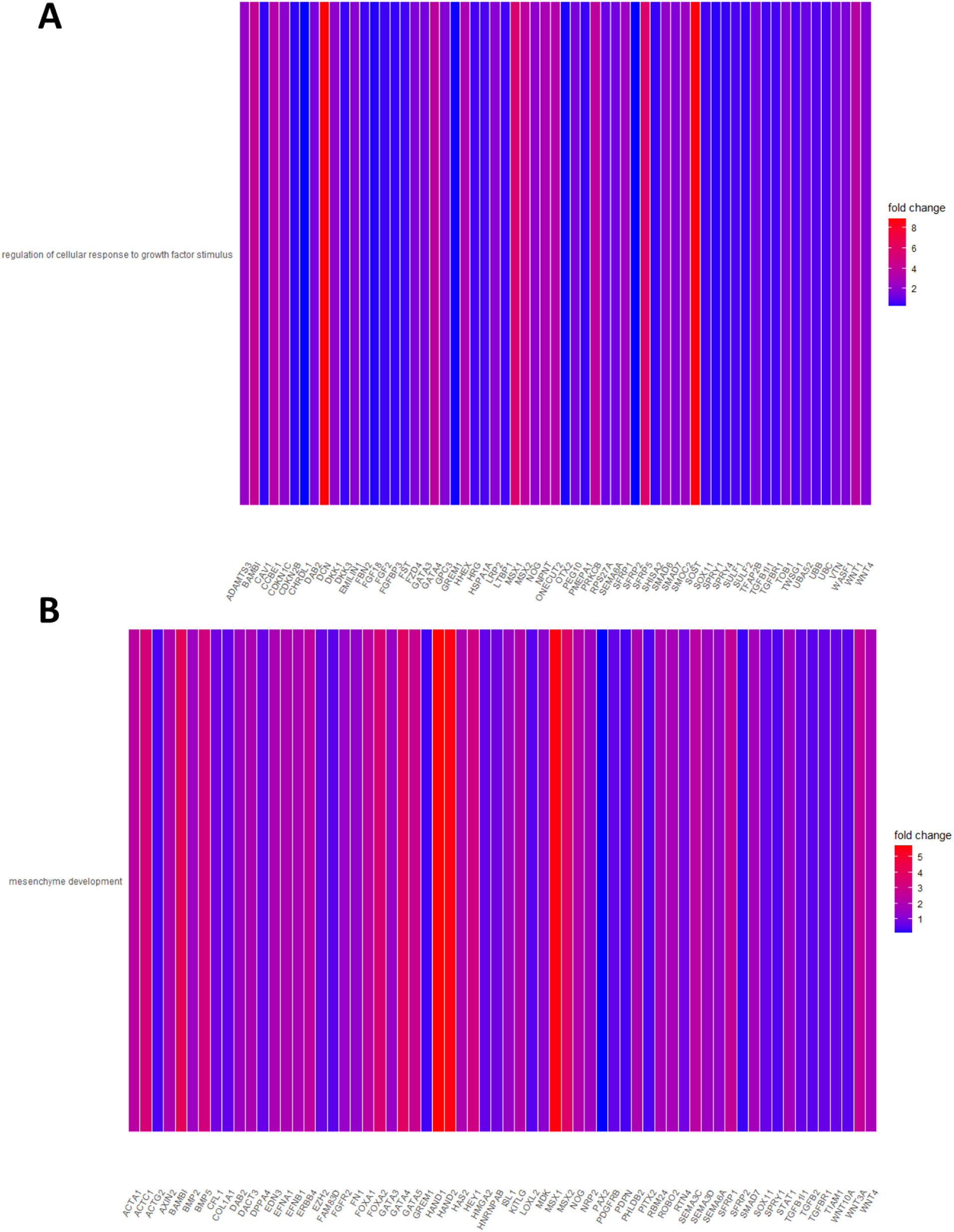
Heatmaps of genes annotated by specific GO terms. A) Heatmap plot showing genes annotated by the GO term: ‘regulation of cellular response to growth factor stimulus’. Cells are coloured by fold change of expression from control group to the experimental group. B) Heatmap plot showing genes annotated by the GO term: ‘mesenchyme development’. Cells are coloured by fold change of expression from control group to the experimental group.

**Supplementary Figure 4).**
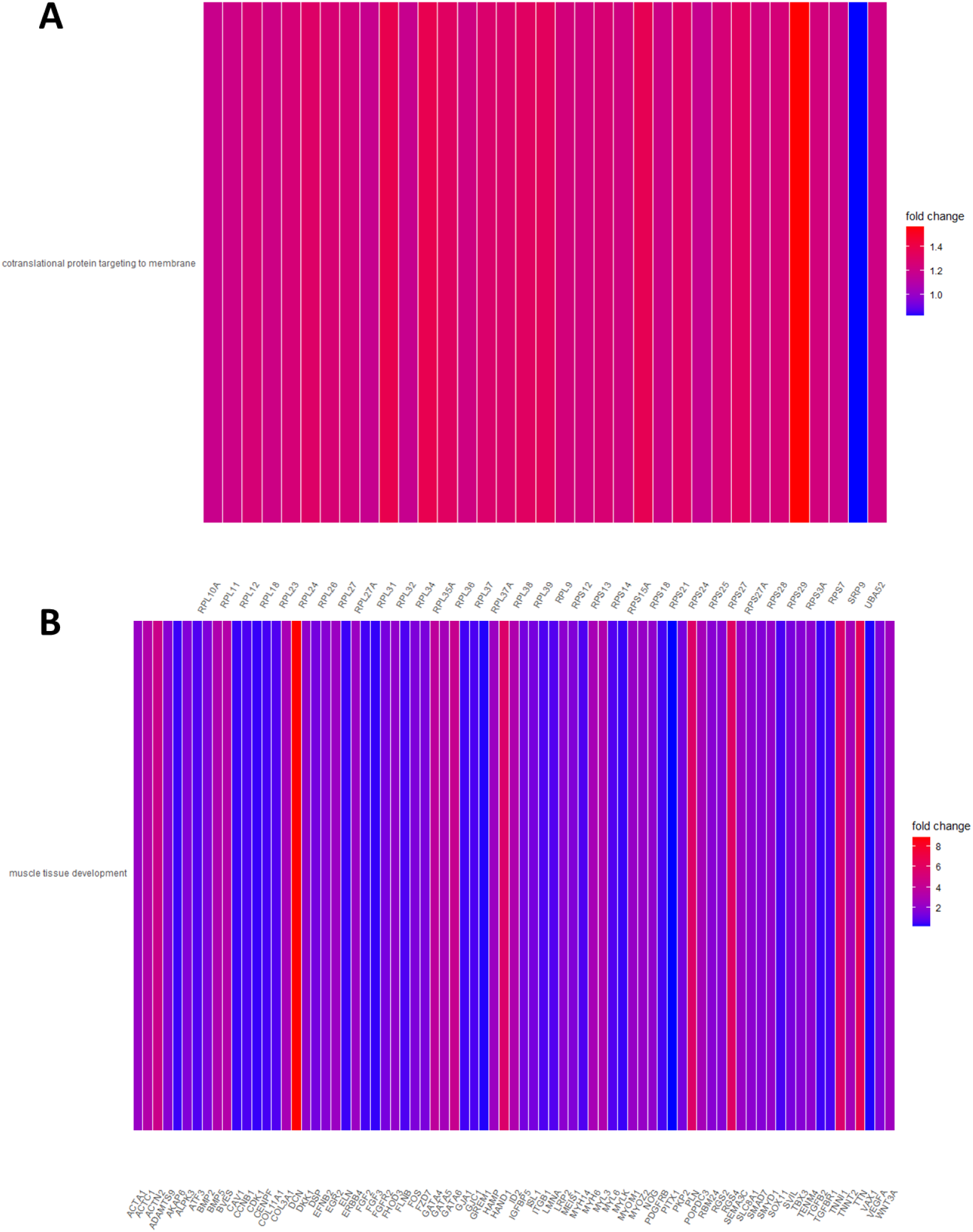
Heatmaps of genes annotated by specific GO terms. A) Heatmap plot showing genes annotated by the GO term: ‘cotranslational protein targeting to membrane’. Cells are coloured by fold change of expression from control group to the experimental group. B) Heatmap plot showing genes annotated by the GO term: ‘muscle tissue development’. Cells are coloured by fold change of expression from control group to the experimental group.

**Supplementary Figure 5).**
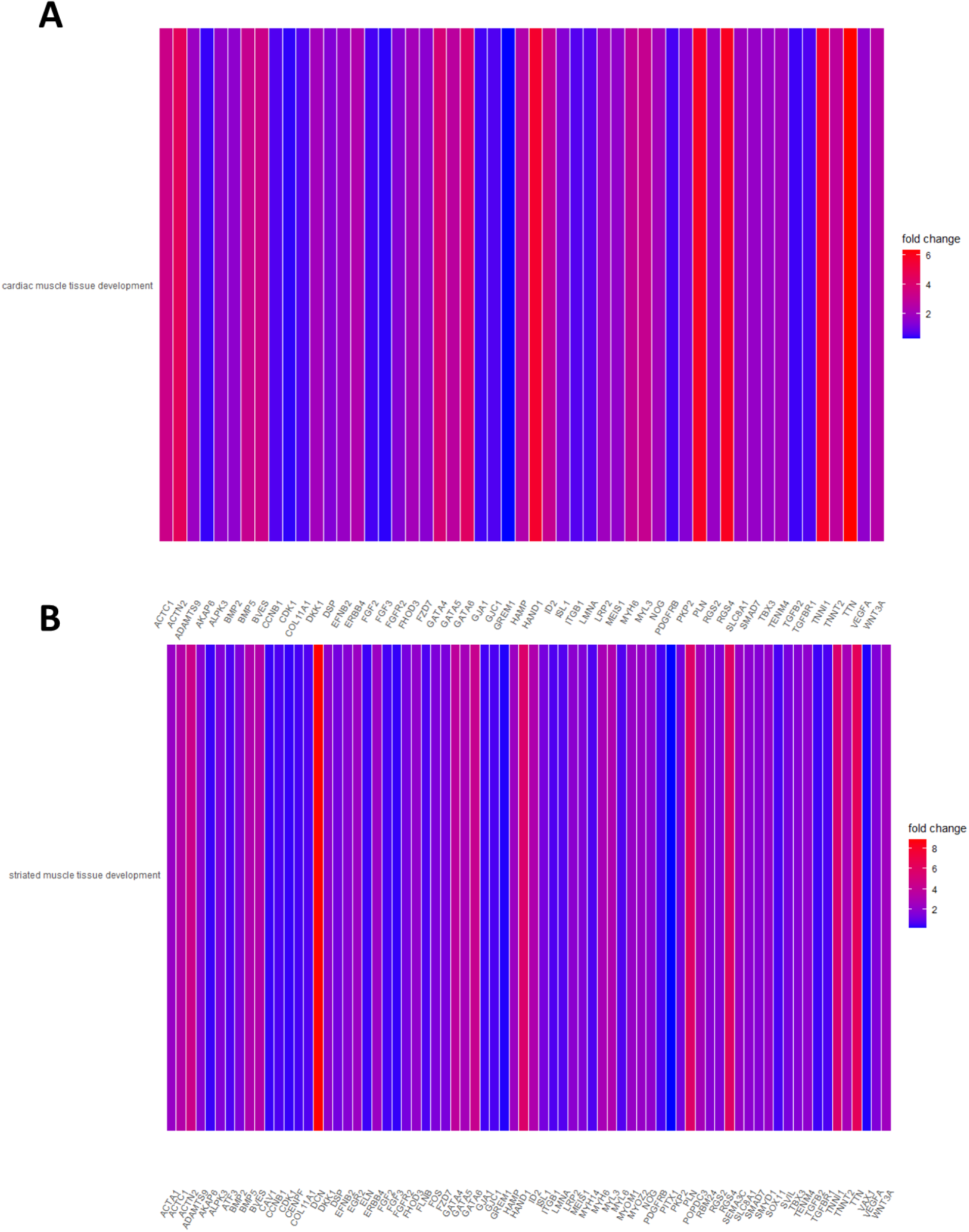
Heatmaps of genes annotated by specific GO terms. A) Heatmap plot showing genes annotated by the GO term: ‘cardiac muscle tissue development’. Cells are coloured by fold change of expression from control group to the experimental group. B) Heatmap plot showing genes annotated by the GO term: ‘striated muscle tissue development’. Cells are coloured by fold change of expression from control group to the experimental group.

**Supplementary Figure 6).**
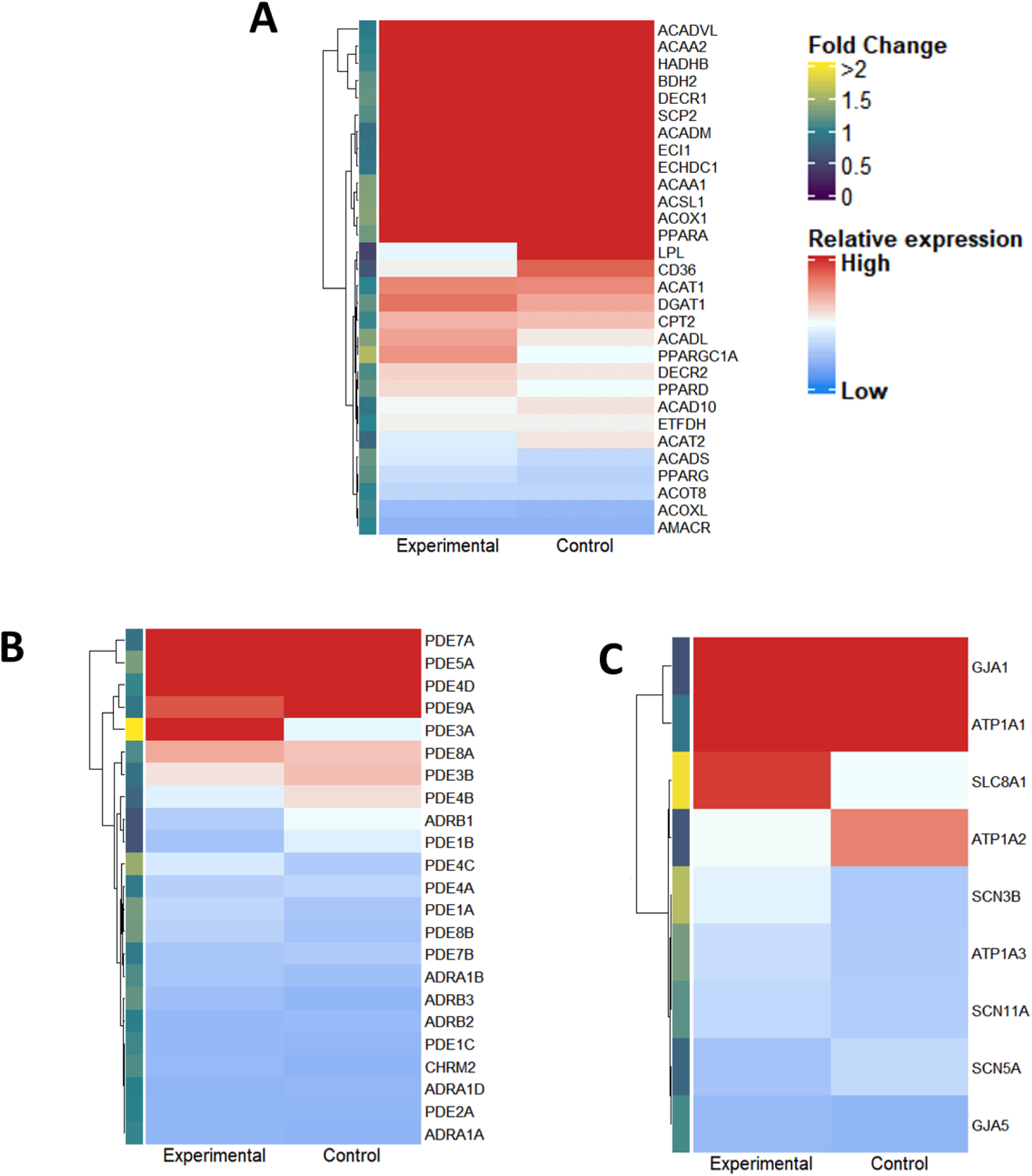
Additional heatmaps of gene expression differences between experimental and control groups across metabolism, signalling, and sodium handling. A) Heatmap plot showing the expression of metabolic genes. Cells are coloured by relative expression. The annotation on the left indicates the specific fold change in expression from the control group to the experimental group. B) Heatmap plot showing the expression of signalling genes. Cells are coloured by relative expression. The annotation on the left indicates the specific fold change in expression from the control group to the experimental group. C) Heatmap plot showing the expression of sodium handling genes. Cells are coloured by relative expression. The annotation on the left indicates the specific fold change in expression from the control group to the experimental group.

## References

1. L.S. Wann, A.B. Curtis, C.T. January, K.A. Ellenbogen, J.E. Lowe, N.A. Estes, 3rd, R.L. Page, M.D. Ezekowitz, D.J. Slotwiner, W.M. Jackman, W.G. Stevenson, C.M. Tracy, V. Fuster, L.E. Rydén, D.S. Cannom, J.Y. Le Heuzey, H.J. Crijns, J.E. Lowe, A.B. Curtis, S. Olsson, K.A. Ellenbogen, E.N. Prystowsky, J.L. Halperin, J.L. Tamargo, G.N. Kay, L. Wann, A.K. Jacobs, J.L. Anderson, N. Albert, J.S. Hochman, C.E. Buller, F.G. Kushner, M.A. Creager, E.M. Ohman, S.M. Ettinger, W.G. Stevenson, R.A. Guyton, L.G. Tarkington, J.L. Halperin, and C.W. Yancy, 2011 ACCF/AHA/HRS focused update on the management of patients with atrial fibrillation (updating the 2006 guideline): a report of the American College of Cardiology Foundation/American Heart Association Task Force on Practice Guidelines. Circulation 123 (2011) 104–23.

2. C.B. de Vos, R. Pisters, R. Nieuwlaat, M.H. Prins, R.G. Tieleman, R.-J.S. Coelen, A.C. van den Heijkant, M.A. Allessie, and H.J.G.M. Crijns, Progression From Paroxysmal to Persistent Atrial Fibrillation: Clinical Correlates and Prognosis. Journal of the American College of Cardiology 55 (2010) 725–731.

3. K. Musunuru, F. Sheikh, R.M. Gupta, S.R. Houser, K.O. Maher, D.J. Milan, A. Terzic, and J.C. Wu, Induced Pluripotent Stem Cells for Cardiovascular Disease Modeling and Precision Medicine: A Scientific Statement From the American Heart Association. Circulation. Genomic and precision medicine 11 (2018) e000043.

4. M. Yazawa, B. Hsueh, X. Jia, A.M. Pasca, J.A. Bernstein, J. Hallmayer, and R.E. Dolmetsch, Using iPS cells to investigate cardiac phenotypes in patients with Timothy Syndrome. Nature 471 (2011) 230–234.

5. I. Itzhaki, L. Maizels, I. Huber, L. Zwi-Dantsis, O. Caspi, A. Winterstern, O. Feldman, A. Gepstein, G. Arbel, H. Hammerman, M. Boulos, and L. Gepstein, Modelling the long QT syndrome with induced pluripotent stem cells. Nature 471 (2011) 225–9.

6. S.D. Lundy, W.Z. Zhu, M. Regnier, and M.A. Laflamme, Structural and functional maturation of cardiomyocytes derived from human pluripotent stem cells. Stem Cells Dev 22 (2013) 1991–2002.

7. S. Yoshida, S. Miyagawa, S. Fukushima, T. Kawamura, N. Kashiyama, F. Ohashi, T. Toyofuku, K. Toda, and Y. Sawa, Maturation of Human Induced Pluripotent Stem Cell-Derived Cardiomyocytes by Soluble Factors from Human Mesenchymal Stem Cells. Mol Ther 26 (2018) 2681–2695.

8. H.D. Devalla, V. Schwach, J.W. Ford, J.T. Milnes, S. El-Haou, C. Jackson, K. Gkatzis, D.A. Elliott, S.M. Chuva de Sousa Lopes, C.L. Mummery, A.O. Verkerk, and R. Passier, Atrial-like cardiomyocytes from human pluripotent stem cells are a robust preclinical model for assessing atrial-selective pharmacology. EMBO Mol Med 7 (2015) 394–410.

9. J.H. Lee, S.I. Protze, Z. Laksman, P.H. Backx, and G.M. Keller, Human Pluripotent Stem Cell-Derived Atrial and Ventricular Cardiomyocytes Develop from Distinct Mesoderm Populations. Cell Stem Cell 21 (2017) 179–194.e4.

10. Z. Laksman, M. Wauchop, E. Lin, S. Protze, J. Lee, W. Yang, F. Izaddoustdar, S. Shafaattalab, L. Gepstein, G.F. Tibbits, G. Keller, and P.H. Backx, Modeling Atrial Fibrillation using Human Embryonic Stem Cell-Derived Atrial Tissue. Sci Rep 7 (2017) 5268.

11. M. Argenziano, E. Lambers, L. Hong, A. Sridhar, M. Zhang, B. Chalazan, A. Menon, E. Savio-Galimberti, J.C. Wu, J. Rehman, and D. Darbar, Electrophysiologic Characterization of Calcium Handling in Human Induced Pluripotent Stem Cell-Derived Atrial Cardiomyocytes. Stem Cell Reports 10 (2018) 1867–1878.

12. L. Cyganek, M. Tiburcy, K. Sekeres, K. Gerstenberg, H. Bohnenberger, C. Lenz, S. Henze, M. Stauske, G. Salinas, W.-H. Zimmermann, G. Hasenfuss, and K. Guan, Deep phenotyping of human induced pluripotent stem cell-derived atrial and ventricular cardiomyocytes. JCI Insight 3 (2018) e99941.

13. M. Lemme, B.M. Ulmer, M.D. Lemoine, A.T.L. Zech, F. Flenner, U. Ravens, H. Reichenspurner, M. Rol-Garcia, G. Smith, A. Hansen, T. Christ, and T. Eschenhagen, Atrial-like Engineered Heart Tissue: An In Vitro Model of the Human Atrium. Stem cell reports 11 (2018) 1378–1390.

14. S. Shafaattalab, E. Lin, E. Christidi, H. Huang, Y. Nartiss, A. Garcia, J. Lee, S. Protze, G. Keller, L. Brunham, G.F. Tibbits, and Z. Laksman, Ibrutinib Displays Atrial-Specific Toxicity in Human Stem Cell-Derived Cardiomyocytes. Stem Cell Reports 12 (2019) 996–1006.

15. Y. Zhao, N. Rafatian, N.T. Feric, B.J. Cox, R. Aschar-Sobbi, E.Y. Wang, P. Aggarwal, B. Zhang, G. Conant, K. Ronaldson-Bouchard, A. Pahnke, S. Protze, J.H. Lee, L. Davenport Huyer, D. Jekic, A. Wickeler, H.E. Naguib, G.M. Keller, G. Vunjak-Novakovic, U. Broeckel, P.H. Backx, and M. Radisic, A Platform for Generation of Chamber-Specific Cardiac Tissues and Disease Modeling. Cell 176 (2019) 913–927.e18.

16. B. Chanda, A. Ditadi, N.N. Iscove, and G. Keller, Retinoic acid signaling is essential for embryonic hematopoietic stem cell development. Cell 155 (2013) 215–27.

17. M. Kawagishi-Hotta, S. Hasegawa, T. Igarashi, Y. Date, Y. Ishii, Y. Inoue, Y. Hasebe, T. Yamada, M. Arima, Y. Iwata, T. Kobayashi, S. Nakata, K. Sugiura, and H. Akamatsu, Increase of gremlin 2 with age in human adipose-derived stromal/stem cells and its inhibitory effect on adipogenesis. Regen Ther 11 (2019) 324–330.

18. I.I. Müller, D.B. Melville, V. Tanwar, W.M. Rybski, A. Mukherjee, M.B. Shoemaker, W.-D. Wang, J.A. Schoenhard, D.M. Roden, D. Darbar, E.W. Knapik, and A.K. Hatzopoulos, Functional modeling in zebrafish demonstrates that the atrial-fibrillation-associated gene GREM2 regulates cardiac laterality, cardiomyocyte differentiation and atrial rhythm. Dis Model Mech 6 (2013) 332–341.

19. V. Tanwar, J.B. Bylund, J. Hu, J. Yan, J.M. Walthall, A. Mukherjee, W.H. Heaton, W.-D. Wang, F. Potet, M. Rai, S. Kupershmidt, E.W. Knapik, and A.K. Hatzopoulos, Gremlin 2 promotes differentiation of embryonic stem cells to atrial fate by activation of the JNK signaling pathway. Stem Cells 32 (2014) 1774–1788.

20. I. Smyrnias, W. Mair, D. Harzheim, S.A. Walker, H.L. Roderick, and M.D. Bootman, Comparison of the T-tubule system in adult rat ventricular and atrial myocytes, and its role in excitation-contraction coupling and inotropic stimulation. Cell Calcium 47 (2010) 210–23.

21. J. Feng, Z. Wang, G.R. Li, and S. Nattel, Effects of class III antiarrhythmic drugs on transient outward and ultra-rapid delayed rectifier currents in human atrial myocytes. Journal of Pharmacology & Experimental Therapeutics 281 (1997) 384–392.

22. D.L. Costantini, E.P. Arruda, P. Agarwal, K.-H. Kim, Y. Zhu, W. Zhu, M. Lebel, C.W. Cheng, C.Y. Park, S.A. Pierce, A. Guerchicoff, G.D. Pollevick, T.Y. Chan, M.G. Kabir, S.H. Cheng, M. Husain, C. Antzelevitch, D. Srivastava, G.J. Gross, C.-c. Hui, P.H. Backx, and B.G. Bruneau, The Homeodomain Transcription Factor Irx5 Establishes the Mouse Cardiac Ventricular Repolarization Gradient. Cell 123 (2005) 347–358.

23. A. Nygren, C. Fiset, L. Firek, J.W. Clark, D.S. Lindblad, R.B. Clark, and W.R. Giles, Mathematical Model of an Adult Human Atrial Cell. Circulation Research 82 (1998) 63–81.

24. B. Goversen, M.A.G. van der Heyden, T.A.B. van Veen, and T.P. de Boer, The immature electrophysiological phenotype of iPSC-CMs still hampers in vitro drug screening: Special focus on IK1. Pharmacology & Therapeutics 183 (2018) 127–136.

25. G. Giannini, A. Conti, S. Mammarella, M. Scrobogna, and V. Sorrentino, The ryanodine receptor/calcium channel genes are widely and differentially expressed in murine brain and peripheral tissues. Journal of Cell Biology 128 (1995) 893–904.

26. S. Iacobas, B. Amuzescu, and D.A. Iacobas, Transcriptomic uniqueness and commonality of the ion channels and transporters in the four heart chambers. Scientific Reports 11 (2021) 2743.

27. N. Niwa, K. Yasui, T. Opthof, H. Takemura, A. Shimizu, M. Horiba, J.-K. Lee, H. Honjo, K. Kamiya, and I. Kodama, Cav3.2 subunit underlies the functional T-type Ca2+ channel in murine hearts during the embryonic period. American Journal of Physiology-Heart and Circulatory Physiology 286 (2004) H2257–H2263.

28. S. Iacobas, B. Amuzescu, and D.A. Iacobas, Transcriptomic uniqueness and commonality of the ion channels and transporters in the four heart chambers. Sci Rep 11 (2021) 2743.

29. E.A. Rog-Zielinska, M.A. Craig, J.R. Manning, R.V. Richardson, G.J. Gowans, D.R. Dunbar, K. Gharbi, C.J. Kenyon, M.C. Holmes, D.G. Hardie, G.L. Smith, and K.E. Chapman, Glucocorticoids promote structural and functional maturation of foetal cardiomyocytes: a role for PGC-1α. Cell Death Differ 22 (2015) 1106–16.

30. D.O. Nelson, P.A. Lalit, M. Biermann, Y.S. Markandeya, D.L. Capes, L. Addesso, G. Patel, T. Han, M.C. John, P.A. Powers, K.M. Downs, T.J. Kamp, and G.E. Lyons, Irx4 Marks a Multipotent, Ventricular-Specific Progenitor Cell. Stem Cells 34 (2016) 2875–2888.

31. S.-p. Wu, C.-M. Cheng, Rainer B. Lanz, T. Wang, Jonathan L. Respress, S. Ather, W. Chen, S.-J. Tsai, Xander H.T. Wehrens, M.-J. Tsai, and Sophia Y. Tsai, Atrial Identity Is Determined by a COUP-TFII Regulatory Network. Developmental Cell 25 (2013) 417–426.

32. E. Grandi, S.V. Pandit, N. Voigt, A.J. Workman, D. Dobrev, J. Jalife, and D.M. Bers, Human atrial action potential and Ca2+ model: sinus rhythm and chronic atrial fibrillation. Circ Res 109 (2011) 1055–66.

33. E. Grandi, S.V. Pandit, N. Voigt, A.J. Workman, D. Dobrev, J. Jalife, and D.M. Bers, Human atrial action potential and Ca2+ model: sinus rhythm and chronic atrial fibrillation. Circulation research 109 (2011) 1055–1066.

34. W.R. Giles, and Y. Imaizumi, Comparison of potassium currents in rabbit atrial and ventricular cells. J Physiol 405 (1988) 123–145.

35. B. Sakmann, A. Noma, and W. Trautwein, Acetylcholine activation of single muscarinic K+ channels in isolated pacemaker cells of the mammalian heart. Nature 303 (1983) 250–3.

36. G. Krapivinsky, E.A. Gordon, K. Wickman, B. Velimirović, L. Krapivinsky, and D.E. Clapham, The G-protein-gated atrial K+ channel IKAch is a heteromultimer of two inwardly rectifying K+-channel proteins. Nature 374 (1995) 135–141.

37. R. Ertl, U. Jahnel, H. Nawrath, E. Carmeliet, and J. Vereecke, Differential electrophysiologic and inotropic effects of phenylephrine in atrial and ventricular heart muscle preparations from rats. Naunyn-Schmiedeberg’s Archives of Pharmacology 344 (1991) 574–581.

38. K. Kikuchi, J.E. Holdway, R.J. Major, N. Blum, R.D. Dahn, G. Begemann, and K.D. Poss, Retinoic acid production by endocardium and epicardium is an injury response essential for zebrafish heart regeneration. Dev Cell 20 (2011) 397–404.

39. L. Cyganek, M. Tiburcy, K. Sekeres, K. Gerstenberg, H. Bohnenberger, C. Lenz, S. Henze, M. Stauske, G. Salinas, W.-H. Zimmermann, G. Hasenfuss, and K. Guan, Deep phenotyping of human induced pluripotent stem cell–derived atrial and ventricular cardiomyocytes. JCI Insight 3 (2018).

40. J.B. Bylund, and A.K. Hatzopoulos, Differentiation of Atrial Cardiomyocytes from Pluripotent Stem Cells Using the BMP Antagonist Grem2. J Vis Exp (2016).

41. F. Liu, D. Long, W. Huang, W. Peng, H. Lan, Y. Zhou, X. Dang, and R. Zhou, The Biphasic Effect of Retinoic Acid Signaling Pathway on the Biased Differentiation of Atrial-like and Sinoatrial Node-like Cells from hiPSC. IJSC 0 (0000).

42. I. Goldfracht, S. Protze, A. Shiti, N. Setter, A. Gruber, N. Shaheen, Y. Nartiss, G. Keller, and L. Gepstein, Generating ring-shaped engineered heart tissues from ventricular and atrial human pluripotent stem cell-derived cardiomyocytes. Nature Communications 11 (2020) 75.

43. M.G. Gunawan, S.S. Sangha, S. Shafaattalab, E. Lin, D.A. Heims-Waldron, V.J. Bezzerides, Z. Laksman, and G.F. Tibbits, Drug screening platform using human induced pluripotent stem cell-derived atrial cardiomyocytes and optical mapping. STEM CELLS Translational Medicine 10 (2021) 68–82.

44. Q. Zhang, J. Jiang, P. Han, Q. Yuan, J. Zhang, X. Zhang, Y. Xu, H. Cao, Q. Meng, L. Chen, T. Tian, X. Wang, P. Li, J. Hescheler, G. Ji, and Y. Ma, Direct differentiation of atrial and ventricular myocytes from human embryonic stem cells by alternating retinoid signals. Cell Research 21 (2011) 579–587.

45. F. Liu, D. Long, W. Huang, W. Peng, H. Lan, Y. Zhou, X. Dang, and R. Zhou, The Biphasic Effect of Retinoic Acid Signaling Pathway on the Biased Differentiation of Atrial-like and Sinoatrial Node-like Cells from hiPSC. IJSC https://doi.org/10.15283/ijsc21148 (2022).

46. S. Koumi, C.E. Arentzen, C.L. Backer, and J.A. Wasserstrom, Alterations in muscarinic K+ channel response to acetylcholine and to G protein-mediated activation in atrial myocytes isolated from failing human hearts. Circulation 90 (1994) 2213–2224.

47. M.A. Skarsfeldt, S.H. Bomholtz, P.R. Lundegaard, A. Lopez-Izquierdo, M. Tristani-Firouzi, and B.H. Bentzen, Atrium-specific ion channels in the zebrafish—A role of IKACh in atrial repolarization. Acta Physiologica 223 (2018) e13049.

48. N. Schmitt, M. Grunnet, and S.-P. Olesen, Cardiac Potassium Channel Subtypes: New Roles in Repolarization and Arrhythmia. Physiological Reviews 94 (2014) 609–653.

49. P. Peterson, M. Kalda, and M. Vendelin, Real-time determination of sarcomere length of a single cardiomyocyte during contraction. American Journal of Physiology-Cell Physiology 304 (2013) C519–C531.

50. G. Iribe, T. Kaneko, Y. Yamaguchi, and K. Naruse, Load dependency in force–length relations in isolated single cardiomyocytes. Progress in Biophysics and Molecular Biology 115 (2014) 103–114.

51. Z. Jian, H. Han, T. Zhang, J. Puglisi, L.T. Izu, J.A. Shaw, E. Onofiok, J.R. Erickson, Y.-J. Chen, B. Horvath, R. Shimkunas, W. Xiao, Y. Li, T. Pan, J. Chan, T. Banyasz, J.C. Tardiff, N. Chiamvimonvat, D.M. Bers, K.S. Lam, and Y. Chen-Izu, Mechanochemotransduction During Cardiomyocyte Contraction Is Mediated by Localized Nitric Oxide Signaling. Science Signaling 7 (2014) ra27–ra27.

52. L. Sala, B.J.v. Meer, L.G.J. Tertoolen, J. Bakkers, M. Bellin, R.P. Davis, C. Denning, M.A.E. Dieben, T. Eschenhagen, E. Giacomelli, C. Grandela, A. Hansen, E.R. Holman, M.R.M. Jongbloed, S.M. Kamel, C.D. Koopman, Q. Lachaud, I. Mannhardt, M.P.H. Mol, D. Mosqueira, V.V. Orlova, R. Passier, M.C. Ribeiro, U. Saleem, G.L. Smith, F.L. Burton, and C.L. Mummery, MUSCLEMOTION. Circulation Research 122 (2018) e5–e16.

53. Y. Wang, F. Yao, L. Wang, Z. Li, Z. Ren, D. Li, M. Zhang, L. Han, S.-q. Wang, B. Zhou, and L. Wang, Single-cell analysis of murine fibroblasts identifies neonatal to adult switching that regulates cardiomyocyte maturation. Nature Communications 11 (2020) 2585.

54. C.Y. Huang, R. Peres Moreno Maia-Joca, C.S. Ong, I. Wilson, D. DiSilvestre, G.F. Tomaselli, and D.H. Reich, Enhancement of human iPSC-derived cardiomyocyte maturation by chemical conditioning in a 3D environment. Journal of Molecular and Cellular Cardiology 138 (2020) 1–11.

55. F.B. Bedada, M. Wheelwright, and J.M. Metzger, Maturation status of sarcomere structure and function in human iPSC-derived cardiac myocytes. Biochimica et Biophysica Acta (BBA) - Molecular Cell Research 1863 (2016) 1829–1838.

56. S. Kannan, M. Farid, B.L. Lin, M. Miyamoto, and C. Kwon, Transcriptomic entropy benchmarks stem cell-derived cardiomyocyte maturation against endogenous tissue at single cell level. PLOS Computational Biology 17 (2021) e1009305.

57. C.W. van den Berg, S. Okawa, S.M. Chuva de Sousa Lopes, L. van Iperen, R. Passier, S.R. Braam, L.G. Tertoolen, A. del Sol, R.P. Davis, and C.L. Mummery, Transcriptome of human foetal heart compared with cardiomyocytes from pluripotent stem cells. Development 142 (2015) 3231–3238.

58. E. Poon, B. Yan, S. Zhang, S. Rushing, W. Keung, L. Ren, D.K. Lieu, L. Geng, C.-W. Kong, J. Wang, H.S. Wong, K.R. Boheler, and R.A. Li, Transcriptome-Guided Functional Analyses Reveal Novel Biological Properties and Regulatory Hierarchy of Human Embryonic Stem Cell-Derived Ventricular Cardiomyocytes Crucial for Maturation. PLOS ONE 8 (2013) e77784.

59. S.M. Kolodziejczyk, L. Wang, K. Balazsi, Y. DeRepentigny, R. Kothary, and L.A. Megeney, MEF2 is upregulated during cardiac hypertrophy and is required for normal post-natal growth of the myocardium. Current Biology 9 (1999) 1203–1206.

60. L. Lai, T.C. Leone, C. Zechner, P.J. Schaeffer, S.M. Kelly, D.P. Flanagan, D.M. Medeiros, A. Kovacs, and D.P. Kelly, Transcriptional coactivators PGC-1alpha and PGC-lbeta control overlapping programs required for perinatal maturation of the heart. Genes Dev 22 (2008) 1948–61.

61. J.J. Lehman, P.M. Barger, A. Kovacs, J.E. Saffitz, D.M. Medeiros, and D.P. Kelly, Peroxisome proliferator-activated receptor gamma coactivator-1 promotes cardiac mitochondrial biogenesis. J Clin Invest 106 (2000) 847–56.

62. E. Mizuta, J. Miake, S. Yano, H. Furuichi, K. Manabe, N. Sasaki, O. Igawa, Y. Hoshikawa, C. Shigemasa, E. Nanba, H. Ninomiya, K. Hidaka, T. Morisaki, F. Tajima, and I. Hisatome, Subtype switching of T-type Ca 2+ channels from Cav3.2 to Cav3.1 during differentiation of embryonic stem cells to cardiac cell lineage. Circ J 69 (2005) 1284–9.

63. P.W. Burridge, E. Matsa, P. Shukla, Z.C. Lin, J.M. Churko, A.D. Ebert, F. Lan, S. Diecke, B. Huber, N.M. Mordwinkin, J.R. Plews, O.J. Abilez, B. Cui, J.D. Gold, and J.C. Wu, Chemically defined generation of human cardiomyocytes. Nature Methods 11 (2014) 855–860.

64. David T.M. Du, N. Hellen, C. Kane, and Cesare M.N. Terracciano, Action Potential Morphology of Human Induced Pluripotent Stem Cell-Derived Cardiomyocytes Does Not Predict Cardiac Chamber Specificity and Is Dependent on Cell Density. Biophysical Journal 108 (2015) 1–4.

65. C. Kane, and C.M.N. Terracciano, Concise Review: Criteria for Chamber-Specific Categorization of Human Cardiac Myocytes Derived from Pluripotent Stem Cells. Stem Cells 35 (2017) 1881–1897.

66. P.J. Reiser, M.A. Portman, X.-H. Ning, and C.S. Moravec, Human cardiac myosin heavy chain isoforms in fetal and failing adult atria and ventricles. American Journal of Physiology-Heart and Circulatory Physiology 280 (2001) H1814–H1820.

67. J.S. Herman, Sagar, and D. Grün, FateID infers cell fate bias in multipotent progenitors from single-cell RNA-seq data. Nature Methods 15 (2018) 379–386.

68. R.A.M. Villanueva, and Z.J. Chen, ggplot2: Elegant Graphics for Data Analysis (2nd ed.). Measurement: Interdisciplinary Research and Perspectives 17 (2019) 160–167.

69. Z. Gu, R. Eils, and M. Schlesner, Complex heatmaps reveal patterns and correlations in multidimensional genomic data. Bioinformatics 32 (2016) 2847–2849.

70. M.I. Love, W. Huber, and S. Anders, Moderated estimation of fold change and dispersion for RNA-seq data with DESeq2. Genome Biology 15 (2014) 550.

71. C. O’Shea, A.P. Holmes, T.Y. Yu, J. Winter, S.P. Wells, J. Correia, B.J. Boukens, J.R. De Groot, G.S. Chu, X. Li, G.A. Ng, P. Kirchhof, L. Fabritz, K. Rajpoot, and D. Pavlovic, ElectroMap: High-throughput open-source software for analysis and mapping of cardiac electrophysiology. Scientific Reports 9 (2019) 1389.

